# A conserved mechanism of membrane fusion in nuclear pore complex assembly

**DOI:** 10.1101/2025.07.21.665908

**Authors:** Jonas S. Fischer, Matthias Wojtynek, Ashutosh Kumar, Harry Baird, Kateřina Radilová, Daria Maslennikova, Kaustubh Ramachandran, Anna N. Becker, Arantxa Agote-Aran, Alessia Loffreda, Annemarie Kralt, Madhav Jagannathan, Gautam Dey, Ulrike Kutay, Stefano Vanni, Karsten Weis

## Abstract

The nuclear pore complex (NPC) forms a large channel that spans the double lipid bilayer of the nuclear envelope and is the central gateway for macromolecular transport between the nucleus and cytoplasm in eukaryotes. NPC biogenesis requires the coordinated assembly of over 500 proteins culminating in the fusion of the inner and outer nuclear membranes. The molecular mechanism of this membrane fusion step that occurs in all eukaryotes is unknown. Here, we elucidate the mechanism by which two paralogous transmembrane proteins, Brl1 and Brr6, mediate membrane fusion in *S. cerevisiae*. Both proteins form multimeric, ring-shaped complexes with membrane remodeling activity. Brl1 is enriched at NPC assembly sites via a nuclear export sequence and then interacts with Brr6 across the nuclear envelope lumen through conserved hydrophobic loops. Disrupting this interaction blocks fusion and halts NPC assembly. Molecular dynamics simulations suggest that the Brl1-Brr6 complex drives membrane fusion by forming a channel across bilayers that enables lipid exchange. Phylogenetic analyses reveal that Brl1/Brr6 homologues are broadly distributed across eukaryotes, and functional experiments in human cells and D. melanogaster establish CLCC1 as an NPC fusogen in metazoans. Together, our results uncover a novel, conserved mechanism for membrane fusion in eukaryotes.

## Introduction

A fundamental challenge for all eukaryotic cells is the selective transport of a diverse set of macromolecules between the nucleus and cytoplasm. This exchange is mediated by the nuclear pore complex (NPC), a ∼50 MDa protein channel that spans the double lipid bilayer of the nuclear envelope (NE). NPCs are among the largest and most conserved cellular complexes in eukaryotes^1–3^. In *S. cerevisiae*, NPCs are composed of over 30 distinct nucleoporins (Nups), which assemble in multiple copies to form a mature structure with eightfold rotational symmetry, totaling more than 500 individual proteins^4–7^.

In recent years, extraordinary progress has been made in characterizing the fully assembled NPC structure, but much less is known about how the large number of NPC subunits come together and how NPCs are inserted into the NE. In general, there are two modes of NPC assembly: post-mitotic assembly, which occurs following NE breakdown during open mitosis, and interphase assembly, in which NPCs are assembled into the intact NE^8,9^. In organisms that undergo closed mitosis, interphase assembly is the only way to build new NPCs^10^, and inserting a functional NPC into the intact NE without compromising nuclear-cytoplasmic integrity presents a great challenge for the cell^11–15^. The stepwise process begins at the inner nuclear membrane, where stable subcomplexes of individual Nups come together to form successive higher order assemblies that culminate in the mature complex^16–19^.

The final step of NPC insertion requires fusion of the inner and outer nuclear membranes to create a continuous pore through the NE^20,16,21^. In *S. cerevisiae,* the two paralogous transmembrane proteins Brl1 and Brr6 are critical for this process^22–28^. Both proteins are essential, and disruption of either halts NPC assembly, resulting in the emergence of unfused, omega-shaped NE herniations that are a hallmark of defective NPC assembly^26,27,29^. Except for Brl1’s ∼280-amino-acid unstructured N-terminus, the two paralogues share a highly similar structure, comprising two transmembrane helices and a luminal amphipathic helix (AH). We and others previously demonstrated that the AH in Brl1 is essential for NPC biogenesis, and single point mutations in the AH stall NPC assembly at a step prior to the fusion of the inner and outer nuclear membranes^27,28^. However, the molecular mechanism of membrane fusion during NPC biogenesis and how Brl1/Brr6 contribute to this process have remained elusive.

In this study, we integrate experimental approaches with molecular dynamics (MD) simulations to uncover how Brl1 and Brr6 mediate membrane fusion during interphase NPC biogenesis. We find that Brl1 and Brr6 interact to form homo-oligomeric ring structures, creating a channel that enables lipid exchange between the inner and outer NE bilayers. Phylogenetic analyses reveal the presence of *BRL1*/*BRR6* homologues across the eukaryotic tree, and functional studies in human cell lines and *D. melanogaster* establish CLCC1 as their functional equivalent in metazoans. Together, our results uncover a novel mechanism for membrane fusion in NPC biogenesis.

## Results

### Brl1 is enriched at NPC assembly sites via a nuclear export signal

Brl1 and Brr6 are paralogues with highly similar domain architecture (**Fig. 1a**). Despite their similarity, both Brl1 and Brr6 are required for NPC assembly in *S. cerevisiae*^23^. Brl1 was found to localize preferentially to the inner nuclear membrane, whereas Brr6 is found in both inner and outer nuclear membranes^26^, and overexpression of one cannot compensate for the loss of the other, suggesting distinct and non-redundant functions^23^. To better characterize the functional differences between Brl1 and Brr6, we focused on the long unstructured N-terminal domain found exclusively in Brl1 (**Fig. 1a**). We first generated a truncated Brl1 variant lacking the N-terminal region (Brl1ΔN) and tested its role *in vivo*. Using a plasmid shuffle strategy (**Fig. S1a**), we replaced the chromosomal *BRL1* and/or *BRR6* copies with wild-type (wt) versions on a counter-selectable plasmid. Then, we stably integrated the *BRL1ΔN* mutant into the genome and selected for cells that lost the wt plasmid. Using this approach, we found that *BRL1ΔN* failed to compensate for a *BRL1* deletion at endogenous expression levels (**Fig. S1b**). However, when overexpressed, Brl1ΔN could functionally replace full-length Brl1. Remarkably, unlike the wt protein, overexpression of Brl1ΔN also rescued the *BRR6* deletion and even restored viability of a *brl1Δ brr6Δ* double deletion strain (**Fig. 1b**). By contrast, overexpression of an N-terminally truncated Brr6 variant could not functionally replace Brl1 but rescued the *BRR6* deletion (**Fig. 1b**). These findings suggest that the N-terminal domain of Brl1 is essential at physiological levels and is important for the functional divergence of Brl1 and Brr6.

**Figure 1:**
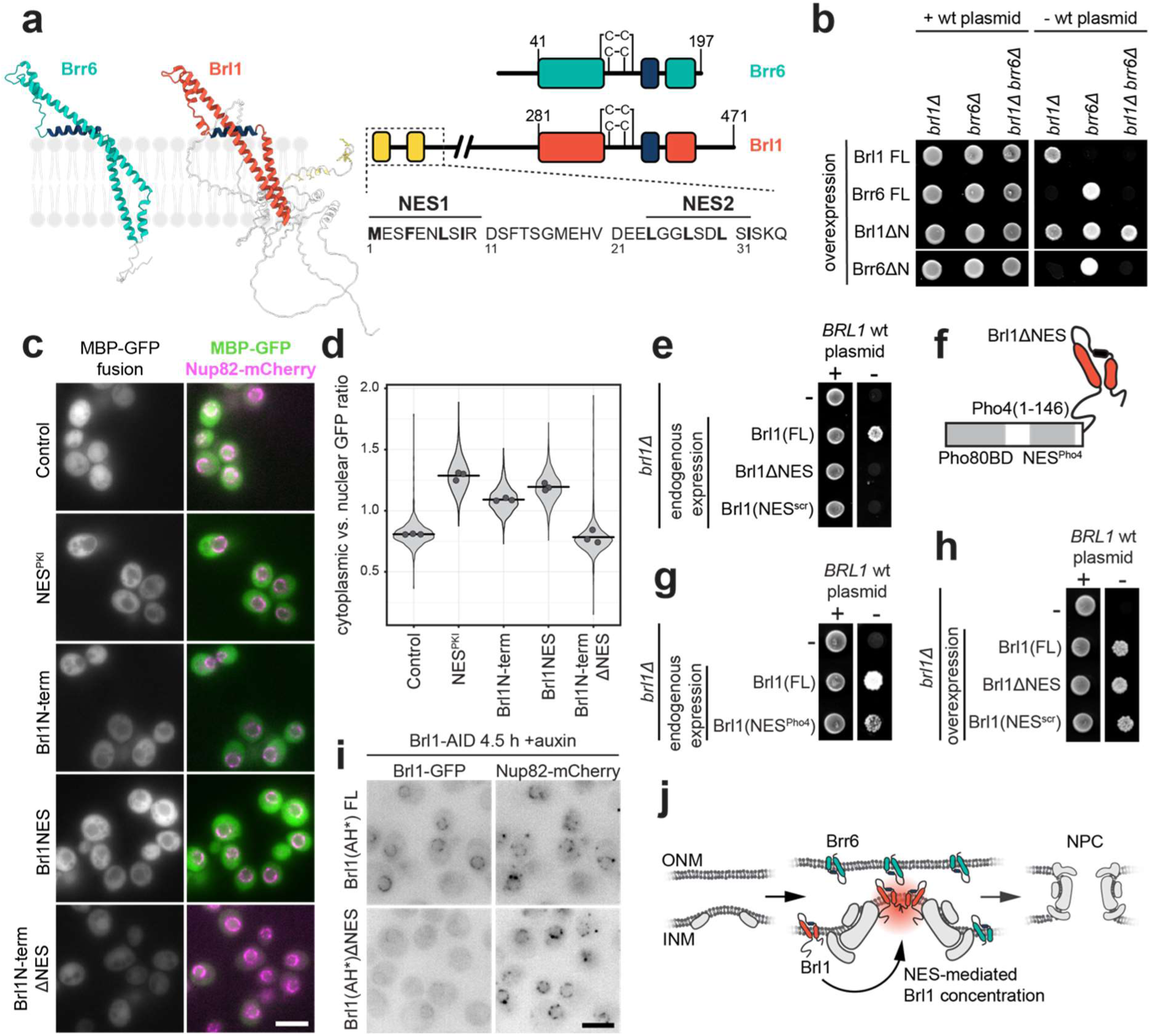
An essential nuclear export signal in the N-terminus of Brl1 is required for its enrichment at NPC assembly intermediates. **a)** AlphaFold3 predictions and domain architecture of *S. cerevisiae* Brl1 and Brr6. Their structured domain is composed of two transmembrane domains (Brr6 teal; Brl1 red), which flank two conserved disulfide bridges and an amphipathic helix (dark blue). Brl1 additionally harbors two predicted Xpo1-dependent nuclear export signals (NES, yellow) in the first 32 residues. These NESs follow the class 1b consensus pattern^91^: Φ-X2-Φ-X2-Φ-X-Φ, where Φ indicates hydrophobic and X any amino acids. **b)** Viability of strains overexpressing either full-length or N-terminally truncated Brl1/Brr6 variants in the presence or absence of their respective wt copies assessed by the plasmid shuffle assay (Fig. S1a). **c)** Representative images of the intracellular localization of N-terminal Brl1 fragments or the *bona fide* NES^PKI^ fused to MBP-GFP. Nup82-mCherry localizes to the nuclear envelope. The control corresponds to MBP-GFP alone. Scale bar: 5 µm. **d)** Quantification of the cytoplasmic to nuclear GFP signal ratio of the strains described in (c). Distribution and mean (points) of three biological replicates are shown. Number of cells per replicate: n ≥ 109 **e)** Viability of strains expressing Brl1 variants from the endogenous promoter in the presence or absence of a plasmid expressing Brl1 wt using the plasmid shuffle assay (Fig. S1a). **f)** Schematic of the Pho4-Brl1ΔNES chimera. A Pho4-fragment including the Pho80 binding site and the Msn5-dependent NES is fused to Brl1ΔNES **g)** Viability of cells expressing the Pho4-Brl1 chimera under the control of the endogenous promoter in the presence or absence of a plasmid expressing Brl1 wt. **h)** Viability of strains overexpressing Brl1 variants in the presence or absence of a plasmid expressing Brl1 wt. **i)** Intracellular localization of the amphipathic helix mutant Brl1(AH*)-GFP with or without the NES. Endogenous wt Brl1 is degraded for 4.5 h using an auxin-inducible-degron (AID). Note that Nup82-mCherry mislocalization indicates an NPC assembly defect. Scale bar: 5 µm. **j)** Model of NES-dependent Brl1 recruitment to NPC assembly intermediates.

Surveying the Brl1 N-terminus using the nuclear export signal (NES) prediction tool LocNES uncovered two potential Xpo1-dependent NESs in the first 32 amino acids of Brl1 (**Fig. 1a**)^30,31^. We tested the functionality of these NESs by fusing N-terminal fragments of Brl1 to GFP-MBP and monitored their intracellular localization. Whereas GFP-MBP alone localized diffusely throughout the cell, constructs containing the full Brl1 N-terminus or just the putative NES fragments were excluded from the nucleus to a similar level as constructs with an NES derived from protein kinase A inhibitor (NES^PKI^)^32^ (**Fig. 1c**). In contrast, constructs lacking the predicted NES, such as GFP-MBP fusions with the Brl1 C-terminus, the Brr6 N-terminus, the Brl1 N-terminus without the NES sequences, or where the amino acids of the NES were scrambled (Brl1(NES^scr^)), localized throughout the cell, similarly to GFP-MBP alone (**Fig. 1c-d, S1c-d**).

Next, we wanted to understand the functional importance of these NES elements in Brl1. Brl1 variants lacking the NES (Brl1ΔNES) or carrying a scrambled version (Brl1(NES^scr^)) were unable to restore viability of *brl1Δ* cells when expressed at low levels (**Fig. 1e**). Importantly, the observed lethality was not due to differences in expression levels, as these constructs were expressed at levels comparable to endogenously tagged Brl1 (**Fig. S1e**). To determine if both NESs are required, we created diploid yeast cells where one copy of *BRL1* was truncated and tested the viability of the haploid cell after sporulation. Interestingly, deletion of either NES alone was tolerated but deletion of both was lethal, indicating that the two NES elements are functionally redundant, but the presence of at least one NES is essential (**Fig. S1f**). Supporting this, fusion of a single, heterologous NES^PKI^ to the Brl1ΔNES mutant could also restore viability (**Fig. S1f**).

Xpo1-dependent NESs are leucine-rich and very hydrophobic in nature^32^. To make sure that it is indeed the nuclear export function that is required and not only the hydrophobicity, we created a chimera between Brl1ΔNES and the longer and less hydrophobic, Msn5-dependent NES from Pho4 (**Fig. 1f**)^33^. A Pho4 fragment containing the NES and one of the two binding domains for the Pho80 kinase was efficiently excluded from the nucleus (**Fig. S1c-d**). Interestingly, fusing this Msn5-dependent NES^Pho4^ to Brl1ΔNES was sufficient to restore the function of Brl1 since its expression rescued the lethality of the *BRL1* deletion strain (**Fig. 1g**). This demonstrates that the presence of an NES in Brl1 is essential, independent of the specific nuclear export pathway.

Brl1 levels play a critical role in NPC assembly, as overexpression of Brl1 has been shown to rescue several NPC assembly mutants such as *nup116Δ* or *gle2Δ*^26,34^. We hypothesized that the use of the export machinery provides a checkpoint mechanism that selectively concentrates Brl1 only at functional NPC assembly intermediates at the inner nuclear membrane. Consistent with this model, overexpression of Brl1ΔNES or Brl1(NES^scr^) restored the viability of *brl1Δ* cells (**Fig. 1h**, compared to **1e**) suggesting that high cellular levels of Brl1 can bypass this enrichment step. To test the NES-dependent enrichment directly, we analyzed the localization of the amphipathic helix mutant of Brl1 (Brl1(AH*)), which is unable to promote NPC assembly, with and without an NES upon degradation of endogenous Brl1 using an auxin-inducible degron (AID)^27,35^. Following auxin-induced degradation of Brl1, full length Brl1(AH*) accumulated in foci around the NE, likely corresponding to stalled NPC assembly intermediates. Brl1(AH*)ΔNES on the other hand was concentrated to a much lower degree (**Fig. 1i**). Furthermore, while overexpression of full-length Brl1(AH*) in a Brl1 wt background is normally dominant negative, cells expressing the truncated Brl1(AH*)ΔNES were viable (**Fig S1g-h**). This suggests that formation of the toxic, multilayered NE herniations we previously observed in this mutant depends on the accumulation of high levels of Brl1(AH*) at the inner nuclear membrane^27^. Together our data reveal a critical role for the NES in localizing and enriching Brl1 at NPC assembly sites. This enrichment mechanism would provide an elegant solution to restrict NPC insertion to assembly sites that can already functionally interact with the nuclear export machinery and prevent erroneous fusion events elsewhere (**Fig. 1j**).

### Brl1/Brr6 are present across the eukaryotic tree and CLCC1 is their functional equivalent in metazoans

In *S. cerevisiae,* both *BRL1* and *BRR6* are essential, and their full-length variants carry out non-redundant functions in NPC biogenesis, at least partly due to the presence of an NES in Brl1. Given that some fungi, such as *S. pombe,* possess only a single *BRL1*/*BRR6* homologue^23^, we sought to investigate the evolutionary ancestry of this protein family. Using HMMER, a tool for sequence-based homology searches^36^, we identified *BRL1*/*BRR6* homologues in 52 of 55 species spanning holomycotan diversity (**Fig. 2a, S2b**). Of note, viability assays revealed that overexpression of orthologues from several fungi could compensate for the loss of Brr6, or even both Brl1 and Brr6, in *S. cerevisiae*, underscoring not only a structural but also remarkable functional conservation of these proteins (**Fig. S2a**). Most fungi contain a single *BRL1*/*BRR6* gene, but a notable duplication appears to have occurred in the Saccharomycotina. Intriguingly, this duplication correlates with the emergence of an N-terminal NES in one of the two paralogues (**Fig. S2b**). This suggests that the gene duplication has enabled the fine-tuning of NPC assembly through the evolution of an NES-mediated targeting mechanism that recruits one Brl1/Brr6 copy to nascent NPCs that form at the inner NE.

**Figure 2:**
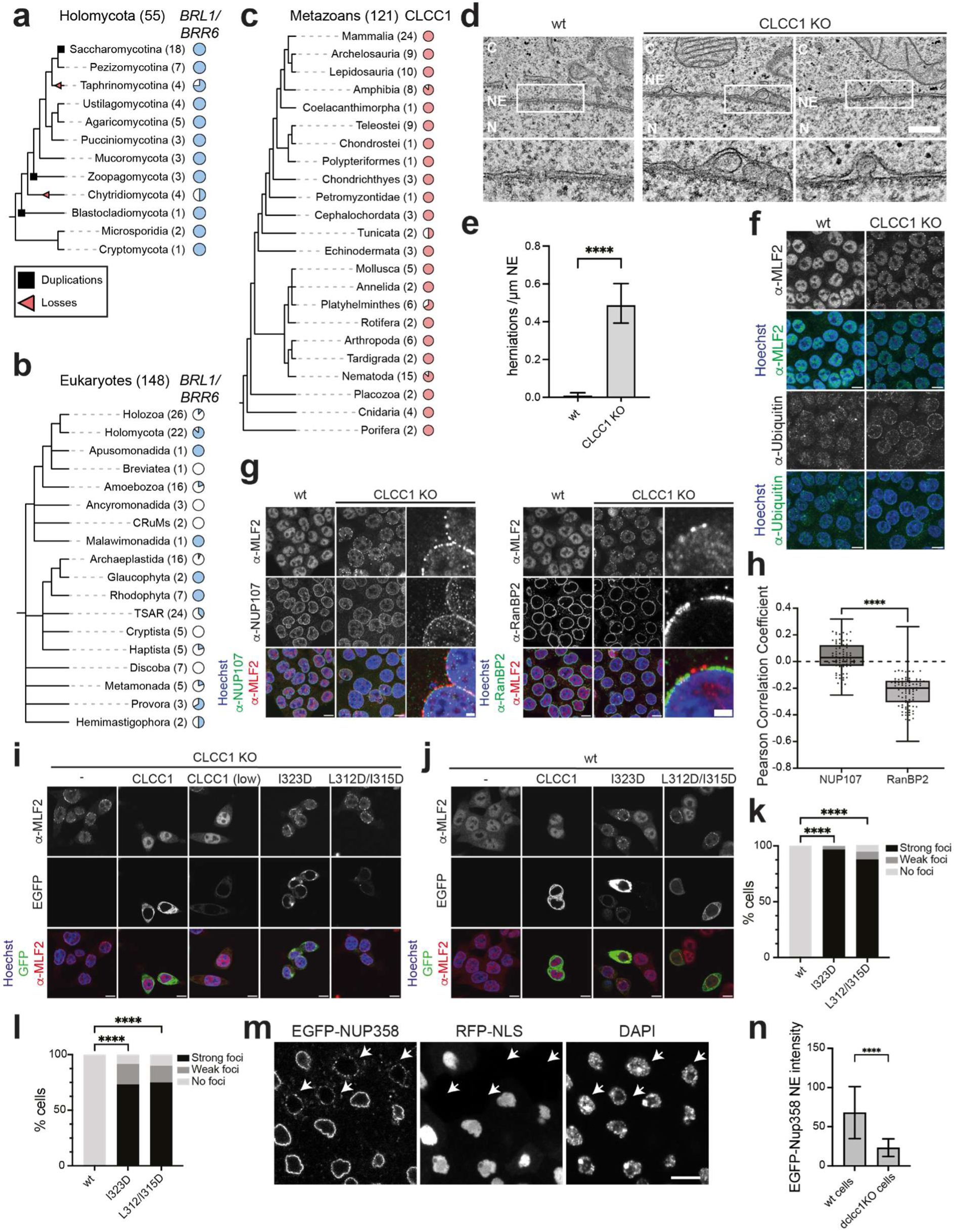
Brl1/Brr6-like proteins are broadly conserved in eukaryotes and CLCC1 is required for NPC biogenesis in human and *Drosophila* cells. **a-c)** Phylogenetic profiles of *BRL1/BRR6* across 55 fungal and other holomycotan species (a), *BRL1*/*BRR6* across 148 eukaryotes (b) or CLCC1 across 121 metazoan species (c). The numbers in the brackets indicate the species selected for each group and the pie charts illustrate the fraction of species for which a homologue was identified. The species trees were manually constructed based on the published literature. In (a), gene duplication or loss events are annotated. Events placed along a branch indicate that they occurred after the divergence of the corresponding lineage. **d)** Representative transmission electron microscopy images of the NE in HeLa wt or *CLCC1* KO cells. N – Nucleus, C – Cytoplasm, NE – Nuclear Envelope. Scale bar: 500 nm. **e)** Quantification of the number of NE herniations in HeLa wt or *CLCC1* KO cells. n = 10 cells; **** p ≤ 0.0001, two-tailed student’s unpaired t-test. Error bars: SD. **f)** Representative images of HCT116 wt and *CLCC1* KO cells stained with anti-MLF2 or anti-ubiquitin antibodies, and Hoechst. Scale bars: 10 µm. **g)** Representative images of HCT116 wt and *CLCC1* KO cells stained with anti-MLF2, and either anti-RanBP2 or anti-NUP107 antibodies, and Hoechst. Scale bars: 10 μm, zoom 2 μm. **h)** Pearson correlation coefficients between the NE signal of MLF2 and either NUP107 or RanBP2 in HCT116 *CLCC1* KO cells. N = 3 biological replicates. Number of cells per condition: n ≥ 80, ****p ≤ 0.0001, Welch’s t-test. Centre line, median; box limits, upper and lower quartile; whiskers, min and max values; points, all values. **i)** MLF2 foci formation in HCT116 *CLCC1* KO cells is rescued by re-expression of CLCC1(wt)-EGFP but not by CLCC1 amphipathic helix mutants. HCT116 *CLCC1* KO cells were transfected with vectors for expression of either CLCC1(wt)-EGFP or CLCC1-EGFP mutants in the amphipathic helix (I323D, L312/I315D). After 48 h, cells were fixed and stained using anti-MLF2 antibody and Hoechst. Scale bars: 10 µm. **j)** Overexpression of CLCC1 carrying mutations in the amphipathic helix causes a dominant-negative phenotype in HCT116 cells. HCT116 wt cells were transfected with vectors for expression of either CLCC1(wt)-EGFP or CLCC1-EGFP mutants (I323D, L312/I315D). After 48 h, cells were fixed and stained using anti-MLF2 antibody and Hoechst. Scale bar: 10 μm. **k)** Quantification of experiment in (i). Cells showing strong, weak or no MLF2 foci at the NE were counted. N = 3 biological replicates, n ≥ 60 per condition. p < 0.001 one-way ANOVA comparing number of “No Foci” cells in each condition. **l)** Quantification of experiment in (j). Cells showing strong, weak or no MLF2 foci were counted. N = 3 biological replicates, n ≥ 60 per condition. p < 0.0001 one-way ANOVA comparing number of “No Foci” cells in each condition. **m)** Cells lacking *dclcc1* (*Drosophila* CLCC1) show defects in NPC biogenesis. dClcc1 is encoded by the CG12945 fly gene. Representative confocal images of EGFP-NUP358 expressing somatic cells of the ejaculatory duct of *D. melanogaster* in a tissue mosaic containing both wt and *dclcc1*^KO^ cells. Cell nuclei are stained with Hoechst. All wt cells express RFP-NLS, while *dclcc1*^KO^ cells lack the RFP signal (arrows). Scale bar: 10 µm. **n)** Quantification of the EGFP-Nup358 signal at the NE from (m). N = 3, n = 60 cells; **** p ≤ 0.0001, student’s unpaired t-test, two-tailed. Error bars: SD. Interactive versions of the phylogenetic trees from a-c) are available at at a) https://itol.embl.de/tree/101163246154401751837272, b) https://itol.embl.de/tree/101163246247831751833282 and c) https://itol.embl.de/tree/1011143160247201751016094

NPC biogenesis is essential in all eukaryotes, yet Brl1/Brr6-like proteins have so far only been reported in fungi, raising questions about their wider evolutionary conservation. To address this conundrum, we extended our search for homologues across 148 species spanning the diversity of eukaryotes. Indeed, Brl1/Brr6-like proteins can be detected throughout the eukaryotic tree (**Fig. 2b**), suggesting an origin predating the divergence of fungi, potentially tracing back to the last eukaryotic common ancestor (LECA). Surprisingly, the distribution of *BRL1*/*BRR6* homologues across the eukaryotic tree is patchy and for many lineages we could not confidently identify homologues. Most notably, no homologue could be found in metazoans using these conventional sequence-based homology searches (**Fig. S3a**). However, HHpred, an algorithm designed to detect distant structural homology^37^, recently identified CLCC1, a metazoan protein with a luminal amphipathic helix flanked by two transmembrane domains, closely resembling the topology of Brl1/Brr6^38–40^. Our phylogenetic analysis showed that CLCC1 is conserved across metazoans (**Fig. 2c, S3b**), but due to high sequence divergence, we were unable to determine if *BRL1*/*BRR6* and CLCC1 share an ancestral origin or if this is a case of convergent evolution. Nevertheless, the structural similarity between these proteins suggests that they may perform equivalent functions in NPC biogenesis.

To test if CLCC1 is required for NPC assembly in metazoa, we created human cell lines in which we deleted CLCC1 by CRISPR/Cas9 (**Fig. S4**) and examined NE morphology using electron microscopy. Consistent with defects in NPC biogenesis, the appearance of NE herniations in a CLCC1 KO cell line, a hallmark of failed NPC assembly^29^, can be observed (**Fig. 2d-e**). Fluorescence microscopy revealed an accumulation of MLF2 and ubiquitin foci at the NE, both established markers of NE herniations in mammalian cells (**Fig. 2f**)^41–43^. To verify that these foci represent stalled NPC assembly intermediates, we assessed the colocalization of MLF2 with NUP107 and RanBP2. NUP107 is incorporated into NPCs prior to membrane fusion, while RanBP2 is recruited only after membrane fusion has occurred^9^. Strikingly, we observed MLF2 foci preferentially in NE regions lacking a RanBP2 signal (**Fig. 2g**). Quantitative analysis indeed revealed that MLF2 and NUP107 signals were uncorrelated as both stalled intermediates and mature NPCs contain NUP107, while there was a clear anticorrelation between MLF2 and RanBP2 localization (**Fig. 2h**). This confirms that the MLF2-positive structures represent pre-fusion NPC assembly intermediates devoid of RanBP2.

To determine whether Brl1 and CLCC1 function similarly in NPC biogenesis, we introduced point mutations in the amphipathic helix of CLCC1 that mimic those previously shown to render Brl1 incapable of promoting membrane fusion^27^. Consistent with its importance for the function of CLCC1, mutations in the CLCC1 AH (I323D or L312/I315D) induced NPC assembly defects, as evidenced by the appearance of MLF2 foci at the NE (**Fig. 2i, 2k**). Like Brl1(AH*) mutants, CLCC1 AH mutants exhibited a dominant-negative phenotype, as defects were observed even in the presence of a wt CLCC1 copy (**Fig. 2j-2l**).

Finally, we examined whether the *Drosophila* CLCC1 homologue *CG12945,* hereafter *dClcc1*, also affects NPC assembly. We obtained a homozygous lethal allele of *dClcc1* (*CG12945^EP3613^, hereafter dclcc1^EP^*) and used the FLP-FRT system to generate mosaic tissues containing *dclcc1^EP^*homozygous clones. In the ejaculatory duct, we observed a marked reduction of Nup358/RanBP2 at the NE in the RFP-null *dclcc1^EP^* homozygous cells, indicating that NPC assembly is halted at a step prior to NE membrane fusion (**Fig. 2m-n**). Taken together, these results demonstrate that CLCC1 is a metazoan-specific NPC assembly factor that not only has a similar structural architecture to Brl1/Brr6 but also functions in the same pathway.

### Brl1 and Brr6 form homo-oligomers with membrane remodeling capability

The broad distribution of Brl1/Brr6 and CLCC1 across the eukaryotic tree, along with their conserved AH, suggests that they may act in a similar manner during NPC assembly. We therefore sought to examine their mechanism of action, focusing on Brl1 and Brr6 in budding yeast. Using AlphaFold3 (AF)^44^, we found that both Brl1 and Brr6 are predicted to self-associate and form curved, multimeric structures that become fully closed rings when 10 or more copies are present (**Fig. 3a, S5a-b**). Notably, similar multimeric ring structures are also predicted for Brl1/Brr6 homologues in other species and for CLCC1, but not for unrelated proteins that also contain two or three transmembrane helices (**Fig. S6-S7**).

**Figure 3:**
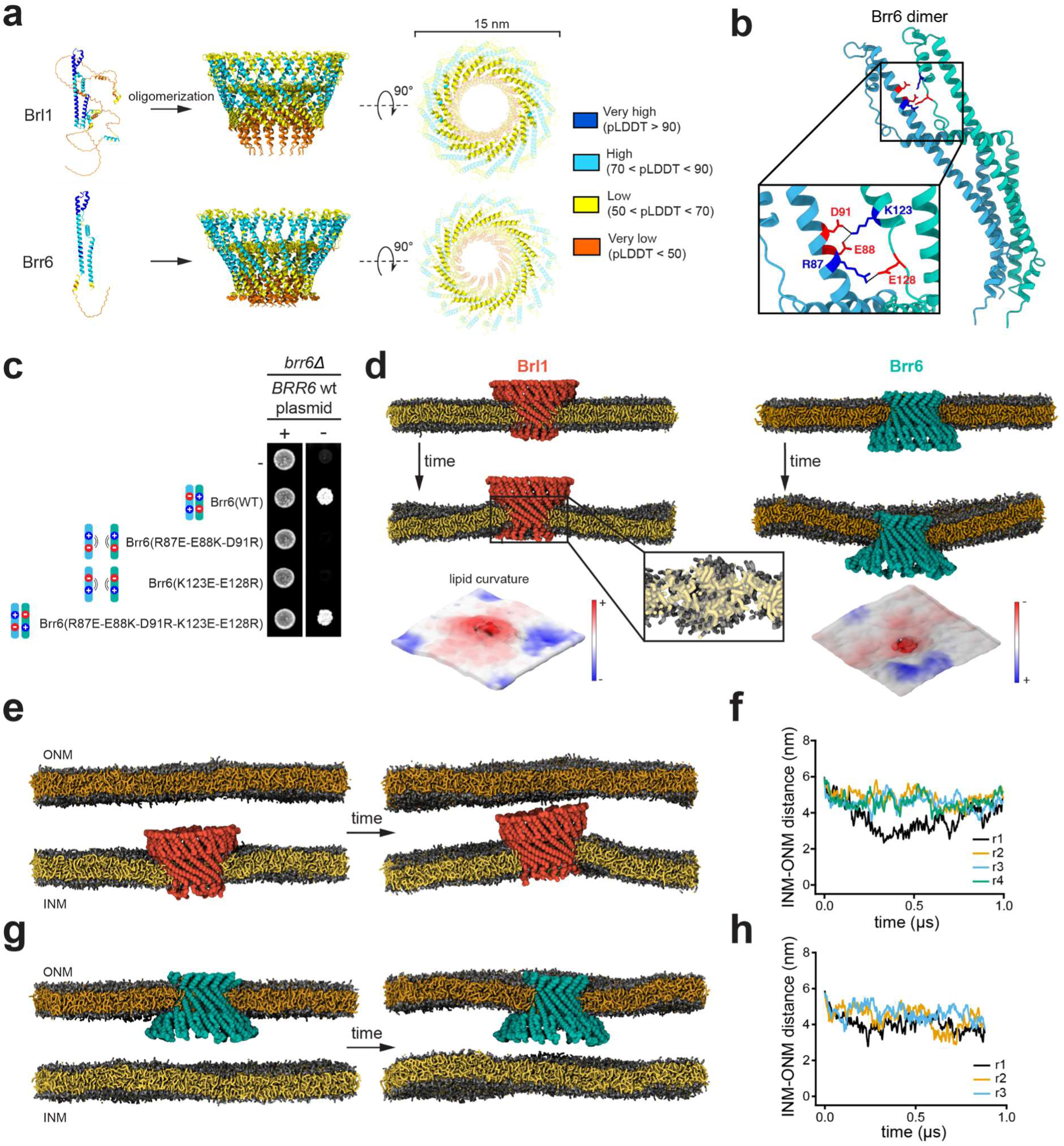
Brl1 and Brr6 form homo-multimeric complexes with membrane remodeling capability. **a)** AlphaFold3 models of Brl1 and Brr6 monomers and homo-oligomers of 16 copies colored by the residue-specific confidence score (pLDDT). **b)** Predicted electrostatic interaction interface between two Brr6 monomers within the 16-mer Brr6 complex. **c)** Viability of cells expressing Brr6 charge-reversal variants using the plasmid shuffling assay. Mutations that introduce electrostatic repulsion between monomers are lethal in the absence of wt Brr6. Reciprocal charge-swapping on both interacting surfaces restores viability. **d)** Coarse-grained MD simulation setup of Brl1 or Brr6 16-mer in a single membrane bilayer along with a final snapshot after the simulation. Inset: Bilayer disruption and lipid remodeling caused by the Brl1 oligomer. Brl1-complex is omitted for clarity and lipid tails are shown as transparent. Bottom: 3-dimensional view of the curvature generation in the membrane leaflets by the Brl1 or Brr6 16-mer. **e)** Coarse-grained MD simulation setup of Brl1 in a double lipid membrane system (left) and a final snapshot after the simulation (right). The spacing between bilayers is 6 nm. **f)** The minimal INM-ONM distance throughout the simulation in (e). **g**) Coarse grained MD simulation setup of Brr6 in a double lipid membrane system (left) and a final snapshot after the simulation (right). The spacing between bilayers is 6 nm. **h**) The minimal INM-ONM distance throughout the simulation in (g). INM – inner nuclear membrane, ONM – outer nuclear membrane.

The AF-predicted protein-protein interactions suggest a conserved oligomerization mechanism in which the amphipathic helices align radially to interact with the membrane (**Fig 3a**). To test this oligomerization model experimentally, we targeted an interaction surface between two Brr6 monomers which is predicted to be mediated by electrostatic interactions (R87, E88, D91 and K123, E128) (**Fig. 3b**). Charge-reversal mutations on either side of this interface disrupted Brr6 function, leading to lethality (**Fig. 3c**). Strikingly, introducing compensatory charge reversals on both sides of the predicted interaction interface restored function and cells expressing this double variant were viable (**Fig. 3c**). This experiment validates the AF model and provides strong evidence that Brr6 multimerization is essential for its function, although the precise stoichiometry of the Brr6 multimer is difficult to determine experimentally. We were unable to engineer equivalent compensatory, electrostatic interaction interfaces in Brl1, but given its structural similarity, we consider it highly likely that Brl1 can also form a multimeric structure as predicted by AF.

To assess the behavior of these oligomeric structures in a membrane environment, we used coarse-grained (CG) MD simulations. To perform the MD simulations of Brl1 and Brr6 complexes inserted into a lipid bilayer, we selected one oligomeric AF model representing 16-mer rings. When embedded in flat lipid bilayers, the oligomeric complexes of both Brl1 and Brr6 introduce pronounced local membrane curvatures and disrupt lipid organization in their vicinity (**Fig. 3d**). To assess this remodeling effect, we calculated the second-rank order parameter (P2) reflecting the degree of order in the lipid tails. We observed a gradual decrease in the order of the lipid tails surrounding the protein assemblies when compared to the bulk membrane region (**Fig. S5d**). Near the oligomers, lipid tails tilt away from their usual orientation, suggesting that they are splayed or inverted, while those farther away remain in a typical bilayer arrangement.

Since such lipid remodeling has been shown to strongly correlate with increased membrane fusion propensity^45^, we wondered whether the observed lipid destabilization might be sufficient to drive membrane fusion. To test this, we performed MD simulations of very closely opposed (6 nm) flat membrane bilayers with a Brl1 or Brr6 16-mer inserted into one of the membranes (**Fig. 3e-h**). However, throughout the simulations the two membranes remained at a constant distance and did not undergo any interaction or fusion (**Fig. 3e-h**). Given that both Brl1 and Brr6 are essential *in vivo*, it was perhaps not surprising that the presence of Brl1 or Brr6 alone was insufficient to induce membrane fusion *in silico*, prompting us to explore how Brl1 and Brr6 cooperate to promote membrane fusion.

### Brl1/Brr6 interact across the ER lumen through a conserved hydrophobic loop motif

Earlier work suggested that Brl1 and Brr6 interact biochemically^25^. To identify their potential interaction interface, we used AF to model assemblies of 16 Brl1 monomers together with 16 Brr6 monomers. Both proteins still formed the previously observed homotypic 16-mer rings and interact with each other side-to-side. Intriguingly, AF predicted that the two homo-oligomers interact in a head-to-head orientation (**Fig. 4a**). This arrangement is consistent with their reported localization in opposing membranes of the NE^26^ and suggests that a luminal interaction between Brl1 and Brr6 might bridge the inner and outer nuclear membranes. The predicted interaction is primarily mediated by short, hydrophobic motifs found at the tips of the luminal domains of both Brl1 and Brr6, composed of four residues: Φ-P-A-Φ (hydrophobic-proline-alanine-hydrophobic), hereafter referred to as the hydrophobic loop (**Fig. 4a**). Multiple sequence alignment revealed that this motif is highly conserved in fungi and other holomycota and likely stabilized by the two previously characterized disulfide bridges^26^, which are also well conserved (**Fig. 4b**).

**Figure 4:**
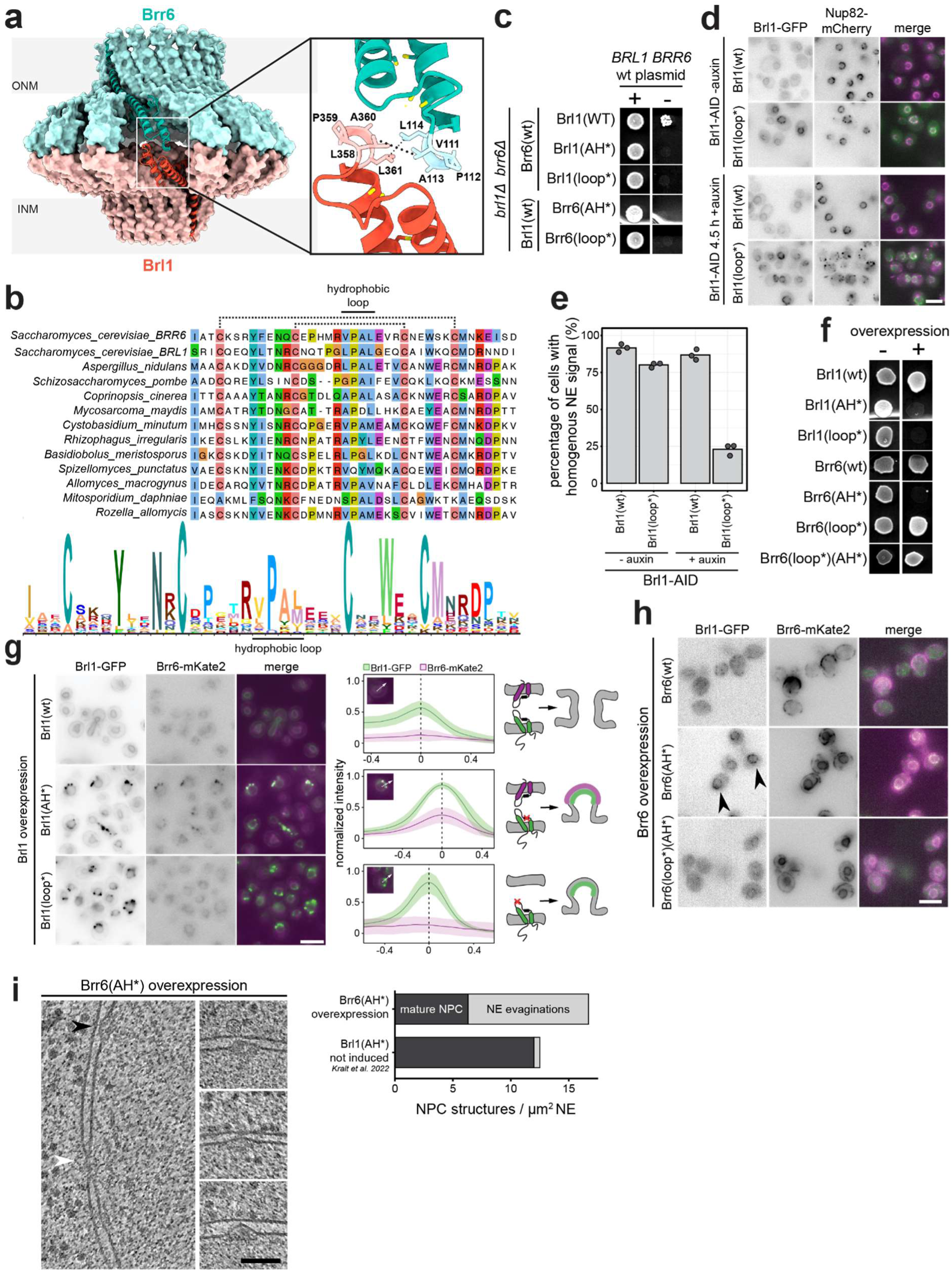
The interaction of Brl1 and Brr6 through conserved hydrophobic loops is essential for NPC assembly. **a)** AlphaFold3-predicted structures of 16 copies of both Brl1 and Brr6. Inset: Predicted head-to-head interaction of Brl1 and Brr6 is localized to a four-residue hydrophobic loop at the tip of their luminal domains. **b**) Top: Multiple sequence alignment of the hydrophobic loop region of *BRL1/BRR6* homologues from the indicated species, spanning the whole phylogenetic profile shown in Fig. 2a. Bottom: The HMM logo diagram of the alignment above. Colored in the CLUSTAL OMEGA color scheme. **c)** Viability assay using the plasmid shuffling of *BRL1* and *BRR6* mutants expressed under the endogenous promoter in the presence or absence of a wt plasmid. **d)** Representative images showing the NPC assembly defects in cells expressing Brl1(wt) or a Brl1 variant, substituting the two leucine residues of the hydrophobic loop by aspartic acids (Brl1(loop*)), upon auxin-induced degradation of endogenous Brl1 (4.5 h, +auxin). Scale bar: 5 µm. **e)** The percentage of cells with a non-homogenous Nup82 NE signal in (d) is quantified as a readout for NPC assembly defects. Points represent the mean of biological replicates. Number of cells per replicate: n ≥ 96. **f)** Viability of cells overexpressing the indicated Brl1 and Brr6 variant in addition to their endogenous copies. **g**) Fluorescence micrographs of cells overexpressing the indicated GFP-tagged Brl1 variants alongside low-levels of Brr6-mKate2 in addition to their endogenous copies. Averaged, normalized intensity profiles across NE accumulations of Brl1-GFP are shown (N = 50 cells). In the Brl1(wt), intensity profiles are drawn across the NE. Scale bar: 5 µm. **h)** Fluorescence microscopy of cells overexpressing Brr6 variants and low-levels of Brl1-GFP in addition to their endogenous copies. Brl1 accumulations are highlighted (black arrow). Scale bar: 5 µm. **i)** Cryo-electron tomography snapshots of cells overexpressing Brr6(AH*), highlighting NPC assembly intermediates (black arrow) and a mature NPC (white arrow). Quantification of NPC structures upon Brr6(AH*) overexpression (n=14) and uninduced *pGAL1-BRL1(AH*)* (n=17) from^27^. Scale bar: 100nm. AH*: mutated amphipathic helix. loop*: mutated hydrophobic loop.

To experimentally test if the hydrophobic loops are indeed required for the interaction between Brl1 and Brr6, we substituted the two hydrophobic residues of the interaction surface with negatively charged aspartic acid (ΦPAΦ->DPAD, subsequently called Brl1(loop*) or Brr6(loop*)). Interestingly, AF no longer predicted a head-to-head interaction between Brl1 and Brr6 for these mutants (**Fig. S8a-b**). Cells expressing loop mutants in either Brl1 or Brr6 were not viable (**Fig. 4c**). Milder substitutions like Brl1(LPAC), Brr6(VPAC) or Brr6(VPAM) were tolerated in the presence of a wt partner, but in combination they resulted in synthetic lethality (**Fig. S8c**), consistent with an interaction between Brl1 and Brr6 via this loop interface.

To monitor how disruption of the hydrophobic loop affects NPC assembly, we employed the AID system to deplete endogenous Brl1 in cells that additionally express either Brl1(wt) or Brl1(loop*). NPC assembly was assessed by the localization of Nup82, a NUP known to incorporate only after membrane fusion has occurred^19,27^. Upon depletion of endogenous Brl1, Brl1(loop*) expressing cells displayed pronounced Nup82 mislocalization, whereas Nup82 remained evenly distributed around the NE in Brl1(wt) cells (**Fig. 4d-e**). These results show that the hydrophobic loop is essential for Brl1 function and that its disruption halts NPC assembly at a step before membrane fusion.

Previously we demonstrated that overexpression of the Brl1(AH*) mutant had a dominant-negative effect, leading to the accumulation of bright foci in the NE and the formation of multilayered membrane herniations^27^. In line with the critical role of the hydrophobic loop in membrane fusion, overexpression of the Brl1(loop*) mutant caused a similar dominant phenotype (**Fig. 4f**). However, fluorescence microscopy revealed a key distinction: whereas the NE foci formed upon Brl1(AH*) overexpression colocalized with Brr6, those formed by Brl1(loop*) did not (**Fig. 4g**). This strongly suggests that although both mutants block NPC assembly, they do so through different mechanisms. Brl1(AH*) retains the ability to interact with Brr6 across the NE lumen but is non-fusogenic because of the disrupted AH. In contrast, Brl1(loop*) fails to interact with and recruit Brr6 to NPC insertion sites, thereby preventing membrane fusion despite having an intact AH.

Overexpression of a Brr6 variant with a mutation in the AH, Brr6(AH*), led to the accumulation of Brl1 in foci at the NE (**Fig. 4h**). This indicates that Brl1 is still recruited to early NPC assembly intermediates and engages with Brr6(AH*), but that Brr6(AH*) is unable to promote membrane fusion, thus halting NPC biogenesis. Indeed, ultrastructural analysis confirmed a substantial increase in arrested NPC assembly intermediates in these cells (10.4 per μm^2^ NE) (**Fig. 4i**) compared to the low frequency observed under normal conditions (0.3-0.5 per µm^2^ NE; ^27^). Like for Brl1(AH*), overexpression of Brr6(AH*) was also dominant-negative (**Fig. 4f**). Remarkably, additional disruption of the hydrophobic loop, Brr6(loop*)(AH*), alleviated the toxicity (**Fig. 4f**) and decreased the accumulation of Brl1 at the NE (**Fig. 4h**). This is consistent with our results that a functional loop is critical for the interaction between Brl1 and Brr6. The Brr6(loop*)(AH*) the double mutant fails to interact with Brl1 and thus permits endogenous Brl1 and Brr6 to function normally in promoting membrane fusion.

In summary, our results show that the hydrophobic loops of Brl1 and Brr6 mediate the luminal interaction between the two proteins. This interaction is essential for membrane fusion, and its disruption on either Brl1 or Brr6 halts the NPC assembly process.

### Brl1-Brr6 interaction leads to lipid mixing between lipid bilayers

Given the importance of the Brl1-Brr6 interaction for NPC assembly, we sought to determine whether the presence of both protein complexes is sufficient to promote membrane fusion. To test this, we performed CG-MD simulations using the AF-predicted structures for the Brl1 and Brr6 homo-oligomeric 16-mers in the 32-mer complex, which featured flatter luminal domains when compared to the individual 16-mers (**Fig 4a, Fig. S5c)**. The two 16-mers were split apart and embedded into opposing membranes (**Fig 5a**). In our MD simulations the two complexes rapidly established stable interactions via the hydrophobic loops (**Fig. 5a-b**) provided that the initial distance between the two bilayers was below 10 nm (**Fig. S9, Supp Movie 3**) and persisted throughout the entire duration of the simulations. Notably, this distance threshold aligns very well with previous membrane-distance measurements of NPC assembly intermediates in cryo-ET tomograms: while the inner and outer NE are normally spaced ∼20 nm apart, they narrow to ∼10 nm as the inner nuclear membrane buckles towards the outer at NPC assembly intermediates^27^. To mimic the high membrane curvature at NPC assembly sites, we also embedded the Brl1 oligomer in a vesicle (R = 12 nm) and again observed robust binding to Brr6 within a few microseconds (**Fig. 5c-d**, **Supp Movie 4**). As a control, no contact between the flat bilayer and the vesicle was established when only an individual 16-mer of Brr6 was present in the bilayer, while leaving the vesicle free of protein. (**Fig. S9**). Overall, this supports our model in which Brl1 and Brr6 interact via their hydrophobic loops and suggests that this interaction can only take place in the context of NPC assembly intermediates.

**Figure 5:**
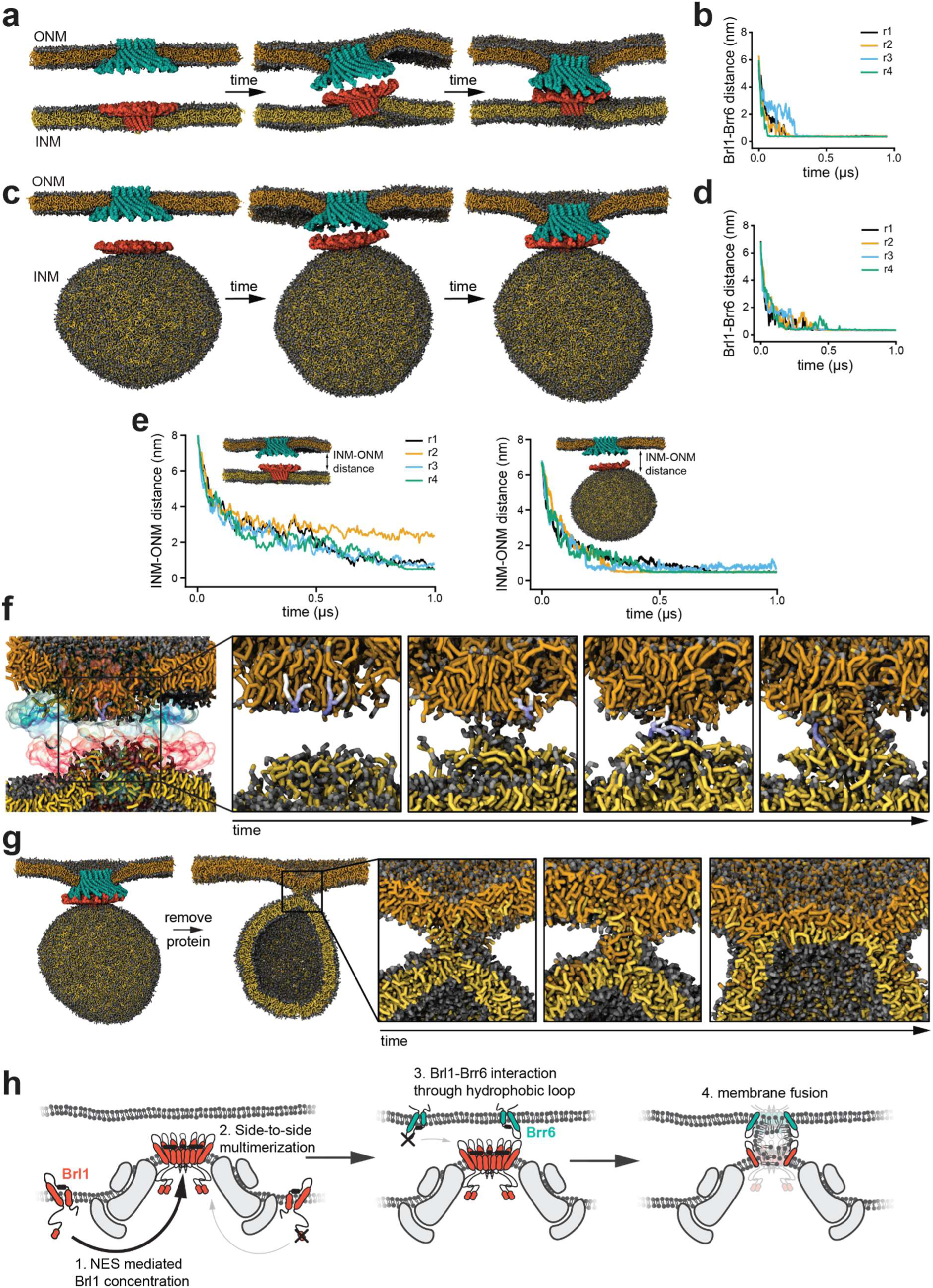
Brl1/Brr6 interaction drives lipid mixing and hemifusion. **a)** Coarse-grained MD simulation featuring two opposing lipid bilayers with Brl1 and Brr6 16-mers inserted into the inner- and outer nuclear membrane, respectively. **b)** Time-trace of minimum Brl1-Brr6 distance during the simulation in (a), showing sustained interaction of the proteins. **c)** Coarse-grained MD simulation of a lipid vesicle containing a Brl1 16-mer and a planar bilayer with a Brr6 16-mer inserted. **d)** Time-trace of minimum Brl1-Brr6 distance during the simulation in (c). **e)** Time-trace of minimum intermembrane (INM-ONM) distance during the simulation in (a and c), indicative of membrane proximity and lipid mixing. **f)** Snapshots illustrating progressive lipid mixing through the interior of the protein oligomeric rings. A blue-marked lipid is tracked to highlight lipid splay prior to mixing. **g)** Transitions from stalk to diaphragm formation in the hemifusion site between two membranes after continuation of the simulation upon protein removal. **h)** Model of nuclear pore complex assembly. INM – inner nuclear membrane, ONM – outer nuclear membrane.

In the MD simulations, the protein-protein interaction between the Brl1 and Brr6 complexes further reduces the distance between the inner and outer NE, effectively pulling the membranes into close proximity to each other (**Fig 5e**). In addition, lipid remodeling similar to that seen in the presence of the Brl1 and Brr6 complexes alone (**Fig. 3d**) was observed on both membranes. Hence, the Brl1-Brr6 complex promotes both membrane tethering and local lipid remodeling near its oligomeric structure, hinting that the formation of this complex primes the inner and outer NE membranes for fusion. Indeed, in all our simulations we observed that once the Brl1-Brr6 interaction is established, lipid mixing takes place within the channel that is formed by the two ring structures (**Fig. 5f**). This lipid mixing is reminiscent of a hemifusion-like intermediate with exposed lipid tails but, in this case, it is stabilized within the protein rings (**Fig. 5f**). We repeatedly found that lipid mixing is exclusively initiated within the channel that is formed by the Brl1-Brr6 ring structure, highlighting the potential role of the oligomeric complex as a catalyst of membrane fusion.

Following lipid mixing, no further spontaneous progression of the fusion process could be observed. To test whether subsequent steps could promote further events in the fusion cascade, we performed additional unbiased CG-MD simulations after removing the protein rings from the final conformation observed in the previous MD simulations with the protein complex. This approach is justified by the observation that Brl1 only transiently associates with assembling NPCs^27^ and that Brl1 and Brr6 are not present in the fully assembled NPC structure^4–6^. Intriguingly, we found that in these conditions, lipid mixing spontaneously progresses into a stable hemifusion stalk intermediate and, over time, all the headgroups rearrange sideways, ultimately facilitating a stalk to diaphragm transition (**Fig. 5g**). Taken together, our simulations identify the hetero-oligomeric Brl1-Brr6 ring complex as a novel catalyst of membrane fusion. Rearrangement or removal of the complex, via a yet unknown mechanism, is likely required to achieve complete fusion between the inner and outer nuclear membrane.

## Discussion

NPCs are a hallmark of all eukaryotic cells, and the insertion of new NPCs into the NE hinges on a pivotal fusion event of the inner and outer nuclear membranes to form a fusion pore that spans the NE. In *S. cerevisiae*, Brl1 and Brr6 have been shown to be essential for this process, yet the underlying mechanism of how Brl1 and Brr6 contribute to membrane fusion has remained unclear. Here, we characterize a heteromeric Brl1-Brr6 ring channel as a novel catalyst of membrane fusion in eukaryotes. Our results demonstrate that Brl1 is targeted via an NES to new NPC assembly sites where it can interact with Brr6 across the lumen of the nuclear envelope. Both Brl1 and Brr6 can multimerize into ring-like homo-oligomers, and upon interaction they can assemble into a large heteromeric fusion channel that bridges the inner and outer NE. MD simulations reveal that the Brl1-Brr6 complex destabilizes lipid bilayers, promoting lipid exchange through the central channel of this ring complex, ultimately driving membrane fusion (**Fig. 5h**).

New NPC assembly is a fundamental process in the life of all eukaryotic cells. Therefore, it has been puzzling that Brl1/Brr6-like proteins have so far only been identified in fungi. Our phylogenetic analyses however demonstrate that homologues of Brl1/Brr6 are present in many lineages across the eukaryotic tree. This suggests that the origin of Brl1/Brr6 predates the divergence of the holomycota clade, and Brl1/Brr6-like proteins might have already been present in the last eukaryotic common ancestor (LECA). In addition, we show that the structurally similar protein CLCC1 is conserved across metazoans and demonstrate that it is required for NPC assembly in human cells and *Drosophila*. Of note, we could not identify Brl1/Brr6 or CLCC1-like proteins in all eukaryotic lineages and, for example, plants lack clearly identifiable homologs. Given the essential nature of NPC assembly, it is likely that there are functionally equivalent proteins in these species as well, but their sequences are too divergent to be identified by sequence-based homology searches. The pronounced sequence divergence between Brl1 and CLCC1 supports the idea that additional, divergent NPC fusogens may exist across eukaryotes. Why such a central component of NPC biogenesis would evolve so rapidly remains unclear. Further investigation into the membrane fusion machinery across diverse eukaryotes will be key to understanding whether this fundamental step is driven by evolutionary conservation or has evolved multiple times and reached a convergent solution.

Despite their elusive evolutionary relationship, our work provides strong evidence that CLCC1 and Brl1/Brr6 perform equivalent roles in NPC assembly. Depletion of human or *Drosophila* CLCC1 halts NPC biogenesis and leads to the accumulation of arrested NPC assembly intermediates (**Fig. 2**). Furthermore, mutations in the AH of hCLCC1 recapitulate the dominant-negative phenotypes previously seen with comparable mutations in the AH of Brl1, suggesting that these two protein families promote membrane fusion in a mechanistically similar way. An obvious difference between these proteins lies in the structure of the hydrophobic loop, which is highly conserved in Brl1/Brr6-like proteins but diverges in CLCC1. Moreover, CLCC1 contains a third transmembrane helix, but can likely also homo-oligomerize into a ring-like structure despite this (**Fig. S6**). While we have identified the AH in CLCC1 as a conserved feature required for membrane fusion, whether CLCC1 mediates fusion through self-interaction or whether it requires additional partners remains an interesting question.

Whereas in most lineages only a single gene coding for Brl1/Brr6 or CLCC1 can be identified, Saccharomycotina have undergone a gene duplication leading to a functional specialization of Brl1 and Brr6, and in *S*. *cerevisiae,* Brl1 and Brr6 can no longer replace each other. We hypothesize that this gives Saccharomycotina better control over the membrane fusion step during NPC assembly. The NES-dependent recruitment of Brl1 that we identify here ensures that the fusion machinery is enriched only at newly forming NPCs that can functionally interact with the nuclear export machinery, thus preventing fusion at non-functional intermediates. Essentially, this could serve as an NPC-assembly checkpoint prior to the initiation of NE fusion. This also predicts that the fidelity of NPC assembly may be lower in our strains that depend on overexpression of Brl1ΔNES or in other eukaryotes that have a single *BRL1*/*BRR6* gene. They likely rely on a homotypic interaction of identical Brl1ΔNES or Brl1/Brr6 rings that could potentially assemble anywhere in the NE or ER. However, insertion of new NPCs and NE fusion must be tightly controlled in all eukaryotes to prevent the uncontrolled exchange of macromolecules between the cytoplasm and the nucleus. Therefore, additional safeguards against uncontrolled NE fusion likely exist. Interestingly, our MD simulations suggest that within the native NE, the two membrane leaflets are too far apart and not flexible enough for the Brl1 and Brr6 rings to interact spontaneously. Therefore, the interaction between Brl1 and Brr6 across the NE lumen can likely only occur at newly emerging NPC assembly intermediates where the initial recruitment of NPC components has already buckled the inner NE membrane and reduced the distance between the inner and outer membranes. Similarly, since Brr6 (and CLCC1) are also found in the ER, it is possible that proteins that control ER luminal width, such as Climp63^46,47^, might play analogous roles to prevent homotopic intraluminal ER fusion. Additional layers of regulation may also contribute. For example, we previously showed that the AH of Brl1 preferentially binds highly curved membranes, such as those found in NPC assembly intermediates^27^. This curvature-sensing mechanism might help to recruit and further stabilize the membrane conformations required for fusion exclusively at new NPC assembly sites.

The observed mechanism of Brl1-Brr6 mediated fusion is conceptually similar to that of other fusion machineries: protein-mediated tethering between two membranes promotes close proximity between the two bilayers, ultimately leading to fusion^48^. Yet, several novel aspects emerge from our data. First, all hitherto known intracellular membrane fusion machineries (e.g., SNAREs, dynamin-like GTPases) promote fusion by cytosolic tethering domains, while the Brl1/Brr6 complex does so using luminal facing domains. Second, SNAREs and dynamin-like GTPases ultimately use energy provided by ATP or GTP hydrolysis to promote membrane fusion but no energy consuming step has so far been identified in yeast NPC biogenesis. It is however important to point out that the energy for membrane fusion in SNARE-mediated processes is provided by tight protein-protein interactions between v- and t-SNAREs that trigger docking and zippering reactions that are sufficient for lipid exchange. ATP hydrolysis is only subsequently required for SNARE complex dissociation and recycling^49^. Akin to SNAREs it appears that the main energetic driving force promoting membrane fusion in NPC biogenesis is the close apposition of the inner and outer NE membrane. This is initially driven by early stages of NPC assembly. Subsequently, the distance between the inner and outer NE is further reduced by the multivalent loop-loop interactions between the Brl1 and Brr6 oligomers, allowing for lipid exchange. How the multimeric complex formed by Brl1 and Brr6 is disassembled and turned over remains an open question. Energy-driven processes, such as post-translational modifications could regulate this late fusion step. Moreover, Brl1/Brr6 together with Apq12 have been implicated in lipid homeostasis^26,50,51^. Thus, a potentially local regulation of the lipid composition within the NE might yet be another important contributor to the fusion process^52,53^. Finally, the post fusion recruitment of additional NPC components, such as the Y-complexes and the cytoplasmic filaments might lead to structural rearrangements within the NPC that stabilize the new fusion pore and might facilitate the recycling of the fusion machinery.

## Acknowledgments

The authors would like to thank members of the Weis lab for discussions and comments on the manuscript, especially Sarah Khawaja and Felix Räsch for their help in revising the manuscript. We acknowledge ScopeM for their support & assistance in this work. We are very grateful to the EMBL IT services HPC resources and team for their assistance. We thank Jana Helsen for her inputs on assembling the set of 55 species to adequately represent the holomycota clade, and her inputs in interpreting the phylogenetic trees for *BRL1*/*BRR6*. We acknowledge ALEMBIC (Advanced Light and Electron Microscopy BioImaging Center) at IRCCS Ospedale San Raffaele for TEM analyses. We thank Cristian Rocha for discussions and inputs on MD simulation setups and Yara Ahmed for help with visualizations.

G.D. and K.Ram. acknowledge EMBL for core funding, as well as the European Union (ERC, KaryodynEVO, 101078291) and FEBS (FEBS Excellence Award to G.D.). K.Ram. holds an Add-on Fellowship for Interdisciplinary Life Science from the Joachim Herz Stiftung. This work was supported by the Swiss National Science Foundation through the National Center of Competence in Research Bio-Inspired Materials and through grants to U.K. (TMAG-3_209245), S.V. (205603) and K.W. (TMAG-3_209354 and 310030_208213).

## Supplementary Information

**Supplementary Video 1 & 2:** Slices through a tomogram used for quantification in Figure 4i in Brr6(AH*) overexpressing yeast strains. Video dimension is 1350nm x 1901nm.

**Supplementary Video 3:** Coarse grained MD simulation of Brl1 and Brr6 16-mers in opposing lipid bilayers.

**Supplementary Video 4:** Coarse grained MD simulation of Brr6 16-mer in a flat lipid bilayer opposed by a Brl1 16-mer in a lipid vesicle.

## Author contribution statement

Conceptualization: K.W., J.S.F., S.V. and U.K.; Investigation and Formal Analysis: J.S.F., M.W., A.Ku., H.B., D.M., A.N.B., K.Rad., K.Ram., A.A.A., A.L. and A.Kr.; Writing – Original Draft, J.S.F., M.W., K.W., S.V., A.Ku.,; Writing – Review and Editing, K.W., J.S.F., S.V., M.W., K.Rad., A.Ku., U.K., M.J., K.Ram., A.A.A., G.D., H.B. and D.M.; Supervision, K.W., S.V., U.K., G.D. and M.J.; Visualization, J.S.F., A.Ku., K.Rad., M.W., H.B., D.M. and K.Ram.; Funding Acquisition, K.W., S.V., U.K and G.D.; Project Administration, K.W.

## Code Availability

Custom code to calculate the surface of the nuclear envelope in this study is available via https://github.com/Mattynek/Brl1project.

## Data Availability

Representative cryo-tomogram have been deposited on wwPDB with the accession code EMD-54440. Strains, plasmids and other data are available from the corresponding authors upon request.

## Materials and Methods

### Cloning procedures

The plasmids for this study were constructed using either scarless homology-based cloning with the In-Fusion® Cloning Kit (Takara Bio) or by modular assembly following the Golden Gate strategy described in^54^. Constructs were either constitutively expressed at low levels under the control of their endogenous promoter or overexpressed using the galactose inducible *GAL1* promoter.

The Brl1(AH*) and Brr6(AH*) mutants correspond to the following point mutations in the amphipathic helix of the two proteins: Brl1(I395D) and Brr6(I149D). In the Brl1(loop*) and Brr6(loop*) mutants, the two hydrophobic residues within a conserved loop region were substituted with aspartic acid: Brl1(LPAL^358–361^ → DPAD) and Brr6(LPAL^111–114^ → DPAD).

The coding region of full-length human CLCC1 was amplified by PCR from human cDNA (forward primer 5’-GATATCATGCTGTGTTCTTTGCTCCTTTG; reverse primer 5’-GCGGCCGCGCCACAGGGGCTGCTG) and cloned into the EcoRV and NotI restriction sites of the pcDNA5.1/FRT/TO vector (Invitrogen), with the EGFP coding sequence inserted between the XhoI and ApaI sites. Brl1 and CLCC1 mutants were generated by gene synthesis (Twist Biosciences).

### CLCC1 CRISPR/Cas9 knockout generation in mammalian cells

HeLa and HCT116 cells were transfected in 6-well plates using JetPrime transfection agent (Polypus), following the manufacturer’s protocol. Each well received 1 µg each of two pC2P-Cas9 plasmids^55^ containing gRNAs targeting the CLCC1 gene: gRNA1 5’-AGTGTTATCTTAACTCTAGA-3’ (targeting ∼400 bp upstream of the start codon) and gRNA2 5’-GTATACTTTACAAAGCTCGA-3’ (targeting ∼40 bp downstream of the stop codon). gRNAs were designed using online platforms CHOPCHOP and CRISPOR^56,57^. To screen for full-gene knockouts, the following primers were used: 5’-CCTAGCAATGGAATTGCTGGG (forward, F1) and 5’-CAAAGAGAAGATGAAATGAAAAGC (reverse, R2). To confirm the presence of the first CLCC1 exon (indicating incomplete knockout), another reverse primer (R1) was used: 5’-GTTGAGAGTTTACTATGCCTCAG. 24 h after transfection, cells were re-seeded onto 10 cm dishes and selected with 2 µg/mL puromycin for two days. After 2.5-3 weeks of regular medium exchange, individual colonies were picked and genotyped.

### Yeast clustering conditions

All yeast strains used in this study were derived from the BY4742 strain (*MAT(alpha) his3Δ1 leu2Δ0 lys2Δ0 ura3Δ0*). Unless otherwise indicated, all yeast strains were cultured to mid log-phase at 30°C in synthetic complete medium (SC, 6.7 g/L yeast nitrogen base without amino acids, 2% sugar) supplemented with the required amino acids and nucleobases. Yeast strains with endogenously tagged protein or proteins under a constitutive promoter were grown in SCD, (SC, 2% dextrose). For the overexpression of Brl1/Brr6 variants, cells were pre-cultured overnight in SC Raff (SC, 2% Raffinose), expression was induced by addition of galactose (2% f.c.) for 6 h. Auxin-inducible degradation of Brl1 was induced by the addition of IP6 (4 μM phytic acid dipotassium salt, Sigma-Aldrich, P5681) and either auxin (+auxin, 500 µM indole-3-acetic acid in ethanol, Sigma-Aldrich, I2886) or the equivalent amount of ethanol (-auxin).

### Cell culture and transient transfection

HeLa and HCT116 cells were cultured in Dulbecco’s Modified Eagle Medium (DMEM; Gibco), supplemented with 10% fetal calf serum (FCS; Eurobio Scientific) and 100 g/ml penicillin/streptomycin (Corning) at 37°C and 5% CO2. For rescue and overexpression experiments, cells were transiently transfected with pcDNA5/FRT/TO (Invitrogen) vectors encoding the GFP-tagged proteins of interest. For each well of a six well plate, a mixture of 100 µl JetPRIME buffer, 2 µl JetPRIME reagent, and 1 µg DNA was prepared and incubated at RT for 15 min before application to the cells. The media was changed after 6 h and the cells fixed after 48 h.

### *Drosophila* strains and husbandry

All fly stocks were raised on standard Bloomington medium at 25°C. The following fly stocks were used: *hs-FLP* (BDSC7), *CG12945^EP3613^*(BDSC17145), *FRT82B* (BDSC86313), *Ubi-NLS-RFP* (BDSC30555) and *EGFP-Nup358* (BDSC95390). To generate *CG12945^EP3613^* homozygous clones, flies were subjected to a 1h heat shock at 37°C approximately 24-48h post egg laying. Somatic tissues (e.g. ejaculatory duct) from adult flies of the genotype *hs-FLP*; *FRT82B, EGFP-Nup358*/+; *Ubi-NLS-RFP*/*FRT82B, CG12945^EP3613^* were dissected and *CG12945^EP3613^* homozygous clones were identified by loss of RFP signal.

### Fluorescent microscopy of yeast cells

Logarithmically growing yeast cells were immobilized in a 384-well glass-bottom plate (MatriPlate) coated with concanavalin A (Sigma-Aldrich). Cells were imaged using a Nikon Ti inverted epifluorescence microscope equipped with a ×100 Plan-Apo VC objective (NA 1.4, Nikon) and a Spectra X LED light source (Lumencore), controlled by NIS Elements software (Nikon). Images were acquired using a Flash 4.0 sCMOS camera (Hamamatsu) and processed with ImageJ.

### Immunofluorescence

Coverslips were fixed in 4% PFA diluted in 1x PBS for 10 min at room temperature (RT), followed by permeabilization with a solution of 0.2% Triton and 0.01% SDS diluted in 1x PBS at RT. Samples were blocked in 2% BSA diluted in 1x PBS for 30 min at RT, incubated with the primary antibodies in 2% BSA-PBS (2 h at RT for -MLF2, -RanBP2, and -NUP107, 4°C overnight for -ubiquitin). Coverslips were washed three times with 2% BSA in 1x PBS at RT, before incubation with secondary antibodies (1:300 in 2% BSA-PBS) in the presence of Hoechst (1 g/ml) for 45 min at RT. The samples were washed three times in 2% BSA-PBS before being post-fixed in 4% PFA in 1x PBS at RT for 3 min. After two washes with 1x PBS, coverslips were mounted with VECTASHIELD (Vector Laboratories) and imaged using a Zeiss 780 upright Confocal Microscope with a 63x 1.4NA Oil Plan-Apochromat objective operated by the ZEN software. Somatic tissues (e.g. ejaculatory ducts) from adult *Drosophila* were dissected in 1x PBS, fixed in 4% paraformaldehyde for 15 min, and washed three times with 0.1% Triton X-100 in 1x PBS for a total of 1 h. Samples were then mounted using VECTASHIELD Antifade Mounting Medium with DAPI (Vector Laboratories) and imaged on a Zeiss LSM 780 upright confocal microscope using a 63×/1.4 NA oil-immersion Plan-Apochromat objective. Mean fluorescence intensity at the nuclear rim was quantified using ImageJ.

### Antibodies

The following antibodies were used for immunofluorescence analysis: mouse monoclonal anti-MLF2 (1:100, Santa Cruz sc-166874), mouse monoclonal anti mono- and poly-ubiquitin conjugates (1:100, Enzo Life Sciences BML-PW8810-500), polyclonal rabbit anti-RanBP2 (1:1000, Abcam ab64276), and rabbit anti-Nup107, raised in rabbits against recombinant protein^58^, anti-mouse Alexa Fluor 488 (Invitrogen), anti-mouse Alexa Fluor 546 (Invitrogen), anti-rabbit Alexa Fluor 488 (Invitrogen). For Western Blot, rabbit polyclonal anti-hCLCC1 (1:500, Atlas Antibodies HPA009087) and mouse monoclonal anti-actin (1:1000, Santa Cruz sc-47778) were used as primary antibodies; mouse Alexa Fluor Plus 680 (Invitrogen, A21058) and rabbit Alexa Fluor Plus 800 (Invitrogen, A32735) as secondary antibodies.

### Quantitative image analysis

To quantify cytoplasmic-to-nuclear fluorescence intensity ratios of the N-terminal Brl1 constructs fused to MBP, segmentation of cells and nuclei was performed using the deep learning-based tool Cellpose (version 3.1.1.1)^59^. For cell segmentation from brightfield images, the following parameters were used: pretrained model cyto2, diameter 75, minimum object size 3, verbose output enabled, exclusion of objects on image edges, no NPY output, cell probability threshold 1, and flow threshold 2. Nuclear segmentation from the Nup82-mCherry fluorescence channel employed the same pretrained model (cyto2) with adjusted parameters: diameter 20, minimum object size 3, cell probability threshold 3.5, and flow threshold 0. Quantification of fluorescence intensities was conducted using CellProfiler4^60^. Cell and nuclear masks generated by Cellpose were imported to define primary ‘Cell’ and ‘Nucleus’ objects. Cells were filtered based on area (1000–10000), form factor (0.7–1), and mean intensity (0.1– 1). Nuclei were filtered by area (100–1000) and form factor (0.7–1). To exclude peripheral regions, filtered cell masks were shrunk by 4 pixels, and cytoplasmic regions were defined by subtracting the nucleus from the shrunken cell mask. To further exclude regions near the plasma membrane and nuclear envelope, both cytoplasmic and nuclear masks were shrunk by an additional 3 pixels. A single manually drawn region per image was used to define background fluorescence. Mean intensities were measured in the shrunken cytoplasmic, nuclear, and background areas. Background-subtracted intensities from cytoplasm and nucleus were used to compute the cytoplasmic-to-nuclear ratio. Only cells with both nuclear and cytoplasmic segmentations were included in the analysis.

Correlation analysis of MLF2 with NUP107 or RanBP2 NE signal was carried out using FIJI (**Fig. 2h**). The NE was defined based on the Hoechst staining and the fluorescence intensities of MLF2 and either NUP107 or RanBP2 was measured along the entire NE. Intensity of the two signals was normalized and the Pearson correlation coefficient was calculated. Only cells showing foci of MLF2 at the NE, and in which the NE remained in a constant z-plane were selected.

Manual, blinded quantification was used to assess the fraction of cells with a homogeneous Nup82-mCherry signal at the NE in cells expressing Brl1(wt) or Brl1(loop*) following Brl1 degradation (**Fig. 4e**).

To quantify co-localization of Brl1 and Brr6 (**Fig 4g**), linescans starting in the nucleus and crossing the nuclear envelope were manually drawn in FIJI for 50 cells per condition. Intensity values were normalized to the maximal intensity, centered to the highest value in the GFP channel and the lowest value of each profile was subtracted. Average and standard deviation were calculated in excel visualized in R.

Brightness and contrast of one fluorescent channel was adjusted the same for all images belonging to one panel using Fiji.

### Transmission Electron Microscopy (TEM)

CLCC1 knockout HeLa cells were fixed in 2.5% glutaraldehyde in 0.1 M cacodylate buffer for 1 h at RT and subsequently washed with the same buffer. The sample was then post-fixed in a solution of 1% osmium tetroxide and 1.5% potassium ferricyanide in 100 mM sodium cacodylate buffer for 1 hour on ice. After several washes in milli-Q water, the sample was incubated in 0.5% uranyl acetate overnight at 4°C, protected from light. Following 5 additional washes with distilled water, the sample was dehydrated in a graded ethanol series (30%, 50%, 70%, 80%, 90%, and 96%, 5 minutes each), followed by 3 washes of 100% ethanol (5 minutes each). The specimen was then infiltrated with a 1:1 mixture of ethanol and epoxy resin and placed on a shaker for 2 hours at RT, followed by two changes of pure epoxy resin (1 hour each) at RT. Finally, the sample was embedded in fresh epoxy resin and polymerized at 45°C overnight, then at 60°C for 24 hours. Ultrathin sectioning (70 nm) were cut using a Leica EM UC7 ultramicrotome and collected on 300 mesh grids. Thin sections were contrasted with uranyl acetate and Sato lead stain. Imaging was performed using a TALOS L120C Transmission Electron Microscope (ThermoFisher Scientific) and images were acquired with a CETA 4x4k CMOS camera (ThermoFisher Scientific).

### Viability assays

To assess the compensation of *brl1Δ, brr6Δ* or *brl1Δ brr6*Δ double deletion mutants with the plasmid shuffling approach, yeast cells were cultured overnight in SCD medium lacking uracil to stationary phase. The following day, cultures were adjusted to an OD₆₀₀ of 2 and spotted onto either SCD -Ura plates or complete SCD plates supplemented with 0.1% 5-Fluoroorotic acid (5-FOA, Zymo Research Corporation, F9003) to select for cells that had lost the *BRL1/BRR6* wt plasmids. For strains overexpressing *BRL1/BRR6* variants under a galactose-inducible promoter, cells were grown overnight in SC Raff -Ura and subsequently spotted onto SC Gal plates. All plates were incubated at 30°C for 3 days and monitored for growth.

### Western blotting

Logarithmically growing yeast cultures equivalent to 2 mL at an OD₆₀₀ of 1 were harvested and lysed by incubation in 1 mL of 0.1 M sodium hydroxide for 15 minutes. Cells were then pelleted and resuspended in 40 µL of Laemmli sample buffer (10% glycerol, 2% SDS, 5% 2-mercaptoethanol, 100 mM DTT, 0.04% bromophenol blue, 62.5 mM Tris–HCl, pH 6.8), followed by heat denaturation at 95 °C for 5 minutes. Proteins were electrophoretically separated by SDS-PAGE using a 10% acrylamide gel and blotted onto a nitrocellulose membrane using the semi-dry Trans-Blot Turbo transfer system (Bio-Rad). Prior to antibody incubation the membrane was blocked for a minimum of 2 h in Immobilon Signal Enhancer (Merck Millipore) at room temperature. Then, the membrane was incubated overnight with primary antibody in Immobilon Signal Enhancer at 4°C, washed 3 x in PBST (PBS, 0.1% v/v tween), followed by incubation with secondary antibody for 1 h at room temperature. Prior to imaging with the Odyssey Imaging System (LI-COR Biosciences), membranes were washed three times for 10 min in PBS. Signal was quantified by dividing the Brl1-GFP signal by the Hxk1 loading control and the resulting ratios from one blot were normalized to endogenous Brl1-GFP signal.

Human cells were lysed in SDS sample buffer (75 mM Tris pH 7.8, 20% (v/v) glycerol, 4% SDS, 50 mM DTT and 0.1% bromophenol blue) and proteins were separated by SDS-PAGE, transferred onto a nitrocellulose membrane (Amersham Protran) via semi-dry blotting. Membranes were blocked in 5% (w/v) milk powder, 0.1% Tween in 1x PBS for 5 min and incubated in primary antibody overnight at 4°C. After three 5 min washes with blocking buffer, membranes were incubated with fluorescent secondary antibodies for 1 hour at room temperature. Following three additional washes in 0.1% Tween-20 in PBS, membranes were imaged using the Odyssey Imaging System.

### Tetrad dissection

Diploid yeast cells were grown on YPD plates for less than 24 h at 30°C and then transferred to sporulation plates (2% agar, 1% potassium acetate, amino acids and nucleotide bases at 25% of regular SCD concentration) and incubated for 5 days at RT. The ascus wall was digested by incubating yeast cells in 5 µL of Zymolyase 100T 1 mg/mL (ICN) for 3 min at 30°C. Then, 300 µL water was added to quench the digestion, cells were briefly vortexed and spread on a YPD plate. Tetrads were dissected using a Nikon Eclipse Ci-S dissecting scope and incubated for 2 days at 30°C. Individual spores were genotyped by sequencing.

### Plunge-freezing

Yeast cells overexpressing Brr6(AH*) were cultured overnight in a selective medium containing Raffinose as the sole carbon source. Cells were pelleted and diluted in the morning to OD 0.6 in medium containing 2% Galactose and incubated at 30°C for 6 hours. Cells were concentrated to OD 2 by spinning and resuspending in Galactose containing medium and plunge frozen on copper grids (Quantifoil R2/1 Cu 200 mesh) using a Vitrobot (Thermo Fisher Scientific). 4ul of cells were blotted from the backside of the grid for 5s using a Teflon sheet on the other side and frozen in a liquid ethane-propane mixture.

### Cryo-FIB milling

Vitrified cells were FIB milled using a Crossbeam 550 (Zeiss) and automated, sequential focused ion beam milling was performed as previously described^61^. In brief, grids were sputter-coated with 4nm Tungsten in an ACE500 (Leica Microsystems) and then transferred to the FIB-SEM where an organometallic platinum precursor was applied using the gas injection system of the machine. FIB milling was performed using currents of 700pA, 300pA, 100pA before fine milling all lamella to a target thickness of 250 nm using 50pA.

### Cryo-electron tomography

Grids containing FIB-milled sections were imaged using a Titan Krios 2 (Thermo Fisher Scientific) at 300kV equipped with a K3 direct electron detector (Gatan) and a GIF BioQuantum energy filter (Gatan). Tilt series were acquired using Tomo5 (Thermo Fisher Scientific) at a magnification of 26,000x and a pixel size of 3.3 Å/pixel. Tilt series were acquired using a dose symmetric scheme with 3° increment, a target defocus of -5 and a cumulative dose of ∼160 e^-^/Å^2^. Tomograms were manually reconstructed in IMOD using patch tracking^62^.

### Quantification of NPCs and NPC evaginations

To calculate the area of NE in Brr6(AH*)-overexpressing cells, we used 13 tomograms with a total NE area of 4.6µm^2^. For comparison with previously published data^27^ we calculated the density of NPC and NPC-like structures per area of NE. For this we manually segmented three slices per NE using the drawing tool in IMOD and interpolated all other slices in between. The model file was then converted using the IMOD function model2point and the area was calculated in MATLAB using these points. NPCs and unfused NPC assembly intermediates were counted manually.

### Mapping the phylogenetic profile of BRL1 in fungi

We selected 55 species, called Holomycota_55, that adequately represent the known diversity of holomycota. We assessed the quality of the selected proteomes using BUSCO v5.7.1^63^ with the eukaryotic lineage of 255 markers (eukaryota_odb10). We identified homologues for the Brl1 protein from *S. cerevisiae* (UniProt ID - P38770) in these species using phmmer from the HMMER set of tools (version 3.4, Aug 2023)^64^. We narrowed down the set of homologues using the best, bidirectional hit approach in which we only retained the best hit from each species that returned P38770 as the top hit in terms of the domain score when used as a query against the *S. cerevisiae* proteome.

We aligned these hits using MAFFTv7.505 using the E-INS-i approach^65^, trimmed the alignments using TrimAl v1.4.rev15 build [2013-12-17]^66^ using the -gappyout option. We redid the trimming using a gap threshold of 0.4 since the -gappyout option had removed over 70% of the positions in the alignment. We removed any sequence at this stage that was composed of over 70% gaps in the trimmed alignment as this could adversely affect the tree building step. We verified the orthology of these sequences using maximum-likelihood (ML) trees built using FastTree (Version 2.1.11 Double precision (No SSE3))^67^. We visualized the trees using FigTree v1.4.4. We used the trimmed alignment to build a Hidden Markov Model (HMM) using hmmbuild from HMMER with the default options. We used this HMM to iteratively search each species in Holomycota_55 separately.

In each iteration, to identify and select new hits in each species, we set an E-value threshold 1E^-4^ for the best one-domain E-value to exclude the least likely homologues. Within the remaining hits, we use the Fisher-Jenks optimization algorithm^68^ to split the hits into two sets based on the best one-domain score such that the variance between the scores in each set is minimized. We then select the set containing the highest score and select all hits in that set. We repeat the alignment-trimming-tree building process as described earlier. We get a new, trimmed alignment that is used to build the HMM for the next iteration of searches. We ran these iterations until no new hits were identified. For the final iteration of sequences, after alignment-trimming as described earlier, we used SMART^69^, PANTHER^70^, SUPERFAMILY^71^, Pfam^72^ analyses from InterProScan 5.73-104.0^73^ to remove sequences that were not annotated to have a “NUCLEUS EXPORT PROTEIN BRR6” (PTHR28136) domain or “Disulfide bridge nucleocytoplasmic transport domain” (PF10104). However, we retained sequences that had no annotation. We built a ML tree using IQ-TREE multicore version 2.3.6 with 1000 ultrafast bootstraps^74,75^. IQ-TREE’s expanded ModelFinder^76^ was used to select the best scoring model as per the Bayesian information criterion. We included one mixture model (LG+C10) in the model search. The final phylogenetic profile was visualized using iToL^77^.

N-terminal NES were predicted using locNES (**Fig. S2b**)^31^. Only the putative NES localized in the first 100 amino acids from the N-terminus with a confidence score > 0.2 were considered. The positions of all predicted NES were checked and NES localized in the transmembrane regions were manually excluded.

### Extending the phylogenetic profile of BRL1 to Eukaryota

We assembled a set of 148 species to represent the known diversity of eukaryotes. We assessed the quality of the proteomes for these species using BUSCO v5.7.1 as described earlier. We used the trimmed alignment generated in the final iteration of the holomycota searches to build a new HMM and individually search each proteome in an iterative manner as described earlier. In each iteration, we further examined the trimmed alignment to verify that at least one pair of cysteines in the [CQEQYLTNRC]NQTP--GLPALGEQ[CAIWKQC] motif, taken from the *S. cerevisiae* BRL1 was present. The phylogenetic profile was generated based off the final alignment and visualized in iToL. We generated the sequence logo using Skylign^78^.

### Mapping the phylogenetic profile of CLCC1 in Metazoa

Similar to the approach described above, we selected a set of 121 metazoan proteomes from UniProt^79^ to represent known metazoan diversity. Since there were no proteomes on UniProt for Ctenophora, we did not include this clade at this stage of the analysis. We used the CLCC1 sequence from *Homo sapiens* (UniProt ID - Q96S66) and followed the same strategy as described earlier to select the best, bidirectional hit for each species and perform iterative HMM searches. We removed sequences without the “CHLORIDE CHANNEL CLIC-LIKE PROTEIN 1” (PTHR34093) annotation or the “Mid-1-related chloride channel (MCLC)” (PF05934) annotation, but we retained those sequences with no annotation. To count the number of CLCC1 copies in each species, we used the UniProt annotations to count the number of unique gene IDs in the final set of CLCC1 paralogues.

### Structure predictions

Multimeric protein structures of Brl1 and Brr6 were predicted using locally-installed AlphaFold2^80^ or the AlphaFold3 web server^44^. The uniprot IDs used are P38770 and P53062 for Brl1 and Brr6, respectively. The quality of the predictions was assessed using the predicted local distance difference test (pLDDT) and interchain predicted template modeling (iPTM).

### MD simulations

All coarse-grained (CG) simulations were performed with GROMACS^81^ v.2023.3, using the Martini 3 force field^82^. To convert the protein structure predictions from all-atom to CG resolution, the Martinize2 script^83^ was used. An elastic network with a force constant of 500 kJ/mol·nm² was applied to preserve the secondary structure of the protein, merging all individual subunits of the protein complex. Next, the insane.py script^84^ was used to generate a lipid bilayer with the following composition: DOPC: DOPE: DOPS: DOPA: DODG (69:20:5:2:4). The bilayer was equilibrated using a five-step protocol that progressively increased the time step while gradually reducing the position restraints on the lipid headgroups, ultimately removing all restraints in the final step to allow the lipids to move freely. The temperature (310 K) and pressure (1 bar) were maintained using the Berendsen thermostat and barostat, respectively. The system was then stripped of solvent and ions to obtain only the equilibrated bilayer which was used in subsequent simulations. For vesicle simulations, we utilized the Martini Maker tool of CHARMM-GUI^85^ to generate a 12 nm-radius vesicle made solely of DOPC. The vesicle was equilibrated following the CHARMM-GUI recommended five-step equilibration approach, with simulation durations of 100, 50, 25, and 25 ns, respectively. The time-steps and position restraints were treated as previously described.

In the first set of systems, containing a single membrane and a protein (**Fig. 3d**), the protein was inserted into the equilibrated bilayer, removing all lipids within 0.7 nm of the protein to prevent steric clashes. Both the protein insertion and the lipid removal steps were executed by an in-house TCL script on VMD^86^. As a result, several lipids remained within the inner ring of the 16-mer oligomers. The system was then solvated with water and ions to achieve charge neutrality and a final excess concentration of 0.15 M NaCl. Energy minimization was performed using the steepest descent algorithm for 5000 steps, with a position restraint of 1000 kJ/mol·nm² applied to the protein backbone beads. The system was then equilibrated using the five-step protocol, incrementally increasing the time step from 1 fs to 15 fs while steadily reducing position restraints on the protein backbone beads. The temperature (310 K) and pressure (1 bar) were maintained using the Berendsen thermostat and barostat, respectively. Four independent replicas of 5 μs were simulated.

For systems with two opposing membranes, we began by building a single membrane containing the desired protein. Each system was built and equilibrated separately using the approach described above. Next, water and ions were removed to obtain the equilibrated structure of the protein in the membrane. The resulting two equilibrated membrane bilayers and protein systems were then positioned apart along the bilayer normal with the minimum desired distance between the opposed membranes. The entire system was then re-solvated with 0.15 M NaCl solution. To maintain the water density in the inter-bilayer space, we implemented an equilibration protocol in which a 2 nm diameter pore was temporarily created in one of the bilayers to allow the passage of water and ions from the intermembrane space to outside of membrane and vice versa. During this process, lipid tails were restrained to stay away from the pore using a flat-bottom restraint potential^87^. The pore was then gradually filled through further equilibration steps by reducing both the diameter and the force constant for the flat bottom restraint, resulting in a stable configuration for production runs. Production simulations across four replicas continued for 1 μs, which was consistently found to be sufficient for lipid mixing to occur between the two bilayers.

To build the systems that contain a protein embedded in a vesicle opposing a protein embedded in a bilayer, we first inserted the protein of interest in the pre-equilibrated vesicle. Following equilibration, we combined the equilibrated protein-in-vesicle system with an equilibrated protein in a single membrane system, ensuring that the proteins were positioned to face each other and that they were at a desired distance from each other. This combined system was then re-equilibrated using a similar five-step protocol. Finally, four independent replicas of these systems were simulated for 1 μs each.

All production simulations were performed with a time step of 20 fs. The temperature was maintained at 310 K using a V-rescale thermostat with a coupling constant of 1 ps, while the pressure was controlled at 1 bar using a Parrinello-Rahman barostat in a semi-isotropic manner, coupling the system laterally while allowing the direction parallel to the bilayer normal to fluctuate freely.

Lipid tail order parameters were computed using the MOSAICS tool^88^. The rank 2 order parameter was computed by taking both tails of lipids into account. To assess the curvature of a membrane throughout the trajectory, SuAVE^89^ was used to project the z-coordinates of all phosphate (PO_4_) beads in the bilayer on a 3D-grid which was visualized as a continuous surface using VMD. For quantification of distances between either two proteins on opposed membranes, or the membranes themselves as a function of time, GROMACS mindist tool was used. For protein-protein minimum distances, all beads of two proteins were considered, while for membranes, only the phosphate (PO_4_) beads of lipids in both inner and outer nuclear membrane were considered for calculation. The distribution of inner-outer nuclear membrane distances was derived from replica-averaged distance profiles. Distances were first averaged across four independent replicas at each time-point, and the resulting distribution was smoothed using a Gaussian kernel with a bandwidth of 0.2.

All graphical plots were generated using Matplotlib and all visual representations were created using either VMD or ChimeraX-1.9^90^.

