## Supplemental Figures for "A conserved mechanism of membrane fusion in nuclear pore complex assembly"

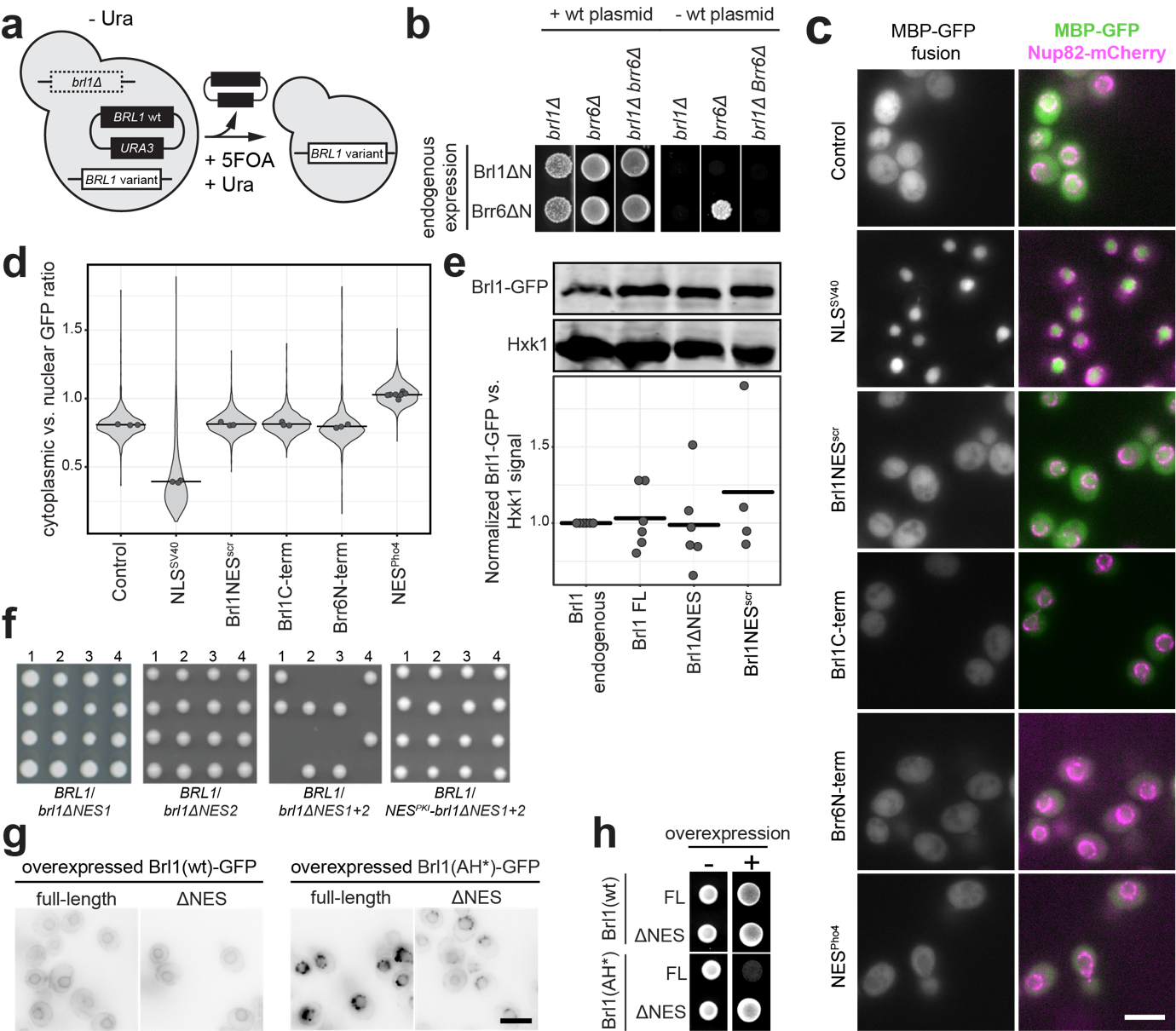

**Supplemental Figure 1: An essential nuclear export signal in the Brl1 N-terminus is required for its enrichment at NPC assembly intermediates. a)** Schematic of the plasmid shuffle assay. Deletion of the endogenous *BRL1* copy is complemented by a *BRL1* wt on a *CEN* plasmid carrying *URA3*. A mutant of *BRL1* is then stably integrated into the chromosome and addition of 5-FOA allows for the selection of cells that lost the *BRL1* wt plasmid during mitosis. **b)** Viability of strains expressing full-length or N-terminally truncated Brl1/Brr6 under their endogenous promoter in the presence or absence of their respective wt copies using the plasmid shuffle assay. **c)** Representative images of the intracellular localization of Brl1/Brr6 fragments, SV40-NLS or the Pho4NES fused to MBP-GFP. Nup82-mCherry localizes to the nuclear envelope. The control corresponds to MBP-GFP alone. Scale bar: 5 µm. **d)** Quantification of the cytoplasmic to nuclear GFP signal ratio of the strains described in (c). Distribution and mean (points) of a minimum of three biological replicates are shown. Number of cells per replicate: n ≥ 127. **e)** Representative western blot and quantification of the mean (bar) of independent replicates (points), comparing endogenously tagged Brl1 to the three Brl1 variants. For the quantification, the Brl1-GFP signals were normalized to Hxk1 and compared to the level of endogenous Brl1-GFP. **f)** Diploid yeast cells carrying one *BRL1* (wt) copy and the indicated *brl1* mutant alleles were sporulated. The resulting tetrads were dissected, and haploid spores were tested for viability. **g)** Representative micrographs of Brl1(wt) or Brl1(AH*) with or without the NES overexpressed for 6.5 h. Scale bar: 5 µm. **h)** Viability assay of yeast cells overexpressing the amphipathic helix mutant Brl1(AH*) with or without an NES.

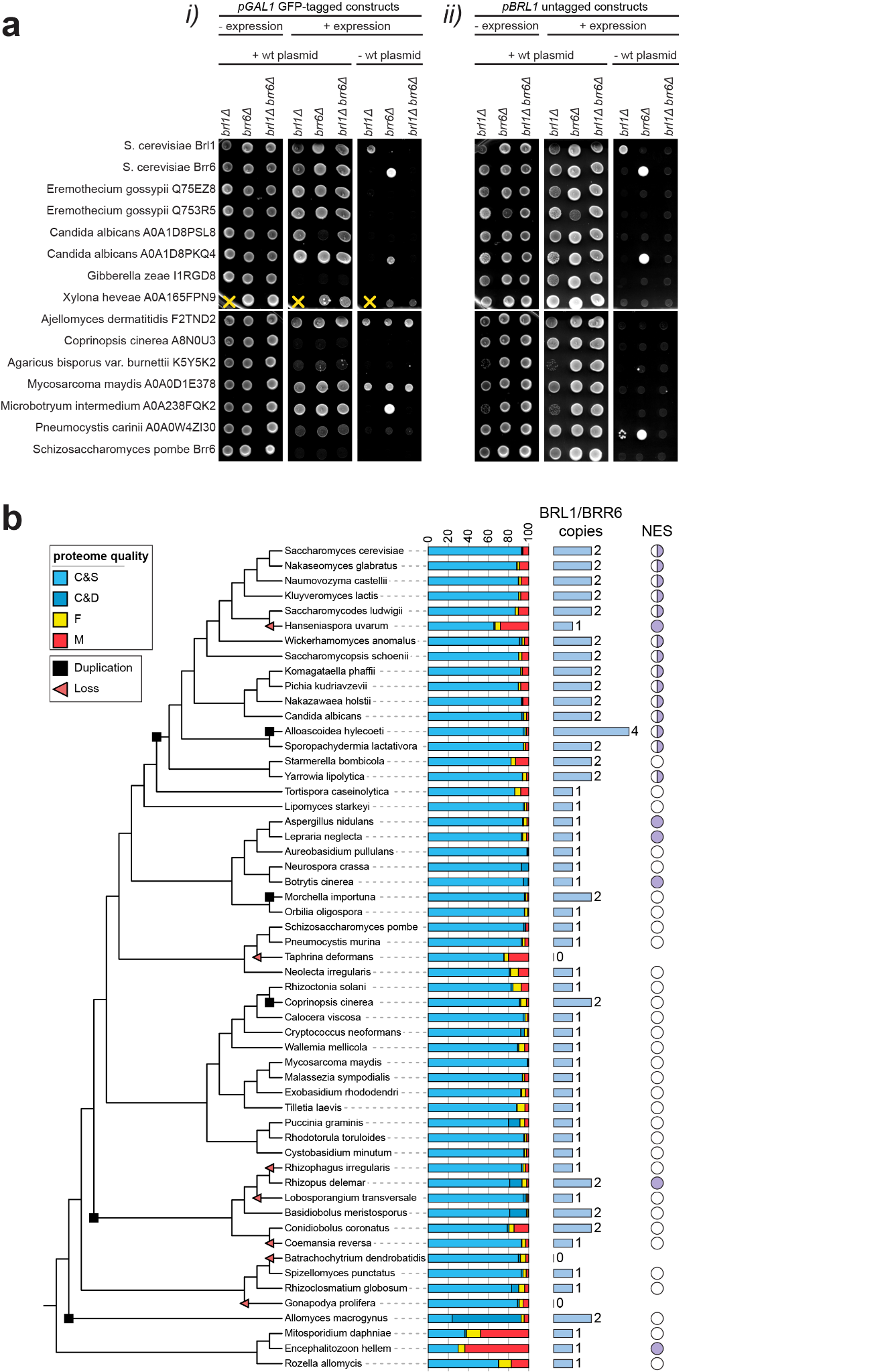

**Supplemental Figure 2: Brl1/Brr6 homologues from diverse fungi rescue the lethality of *brl1Δ brr6Δ* in *S. cerevisiae.*** **a)** Viability of *brl1∆*, *brr6∆* or *brl1∆* *brr6∆* double deletion strains expressing *BRL1/BRR6* homologues from various fungal species. Homologues were expressed either *(i)* at high levels under the control of the *pGal1* promoter and tagged with GFP, or *(ii)* at low levels under the control of the native *pBRL1* promoter without a tag. All constructs were codon-optimized for expression in *S. cerevisiae.* Yellow crosses denote strains that were not tested. **b)** Full phylogenetic profile of *BRL1*/*BRR6* copy numbers across the 55 fungal and other holomycotan species from Fig. 2a. The species tree is manually constructed based on published literature. Proteome quality was assessed using BUSCO v5.7.1. C&S - Completed and Single score, C&D - Completed and Duplicated score, F - Fragmented score, M - Missing score. The duplication/loss events were annotated in the tree based on the distribution of *BRL1* copy numbers. The annotation was based on minimizing the number of evolutionary events needed to account for the extant copy number. The NES column indicates the fraction of proteins for which locNES^31^ predicts an NES in the first 100 N-terminal residues. An interactive version of this phylogenetic tree is available at: <https://itol.embl.de/tree/101113541411731746628869>

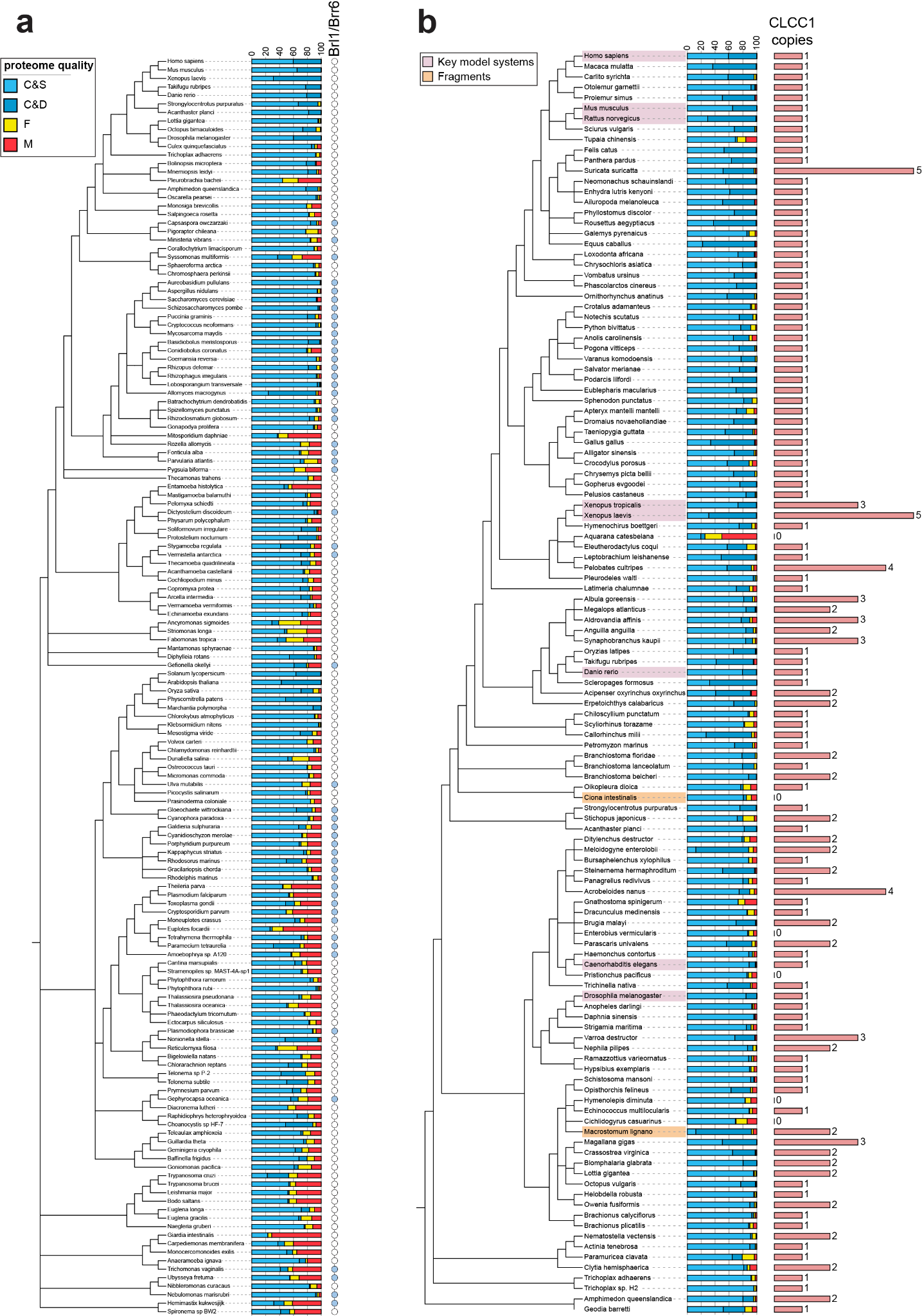

**Supplemental Figure S3: Complete phylogenetic profiles from Fig. 2b-c. a)** Full phylogenetic profile of *BRL1*/*BRR6* homologues across 148 eukaryote species from Fig. 2b. A blue circle indicates that a *BRL1*/*BRR6* homologue was found. An interactive version of this phylogenetic tree is available at: <https://itol.embl.de/tree/1011131123130351748421541> **b)**  Full phylogenetic profile of CLCC1 across 121 metazoan species from Fig. 2c. The bar indicates the number of identified CLCC1 homologues. The species trees were manually constructed based on published literature. Proteome quality was assessed using BUSCO v5.7.1. C&S - Completed and Single score, C&D - Completed and Duplicated score, F - Fragmented score, M - Missing score. An interactive version of this phylogenetic tree is available at: <https://itol.embl.de/tree/1011143160247201751016094>

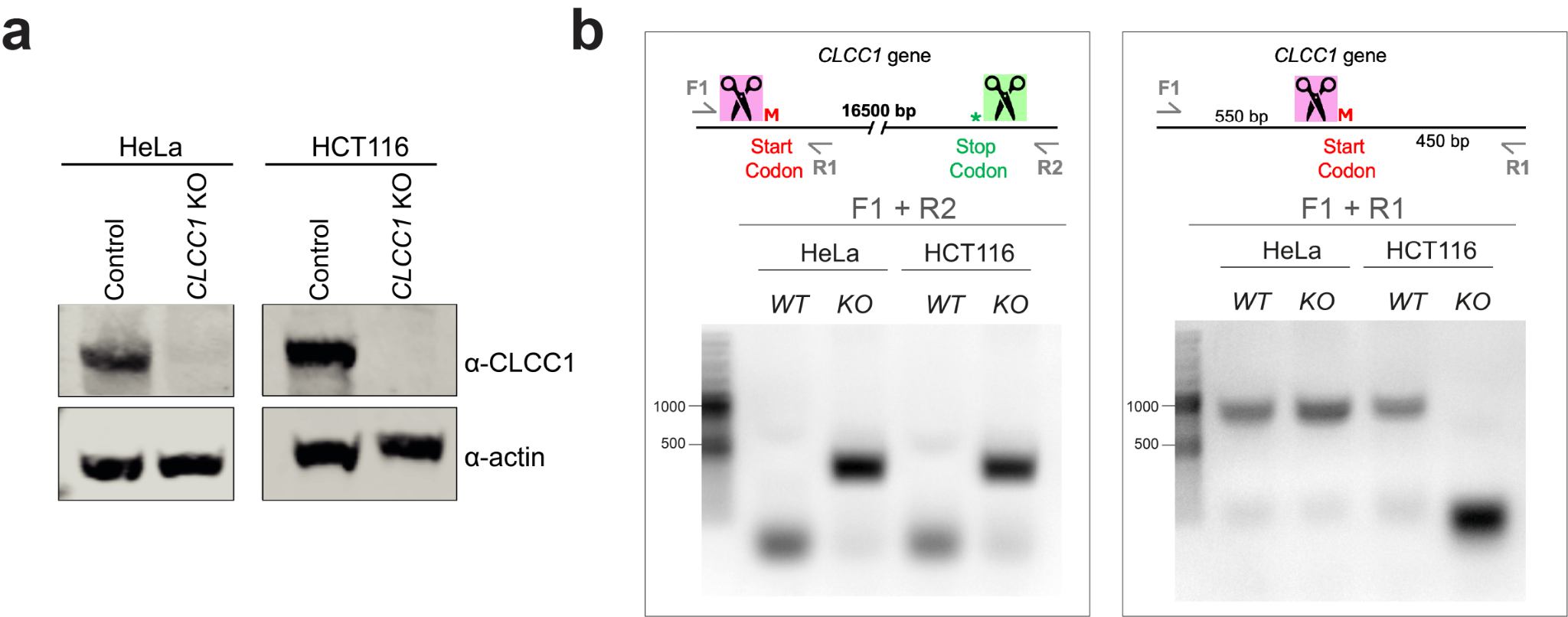

**Supplemental Figure S4: Characterization of CLCC1 knockout generation. a)** Immunoblots confirming deletion of the CLCC1 gene using CRISPR/Cas9 in HeLa and HCT116 cells. Note that a faint residual CLCC1 protein band in the HeLa sample indicates incomplete knockout. **b)** PCR analysis was performed to confirm the deletion of the CLCC1 gene. A forward primer (F1) located upstream of the start codon (and the 5' guide RNA cutting site shown in pink) and a reverse primer (R2) downstream of the stop codon (and the 3' guide cutting site shown in green) were used to confirm knockout of all coding exons. In wt alleles, this region is too large to be amplified (~16 kbp), while upon successful deletion of the gene, a PCR product of ~400 bp is obtained (shown for both HeLa and HCT116 cells). An additional primer pair, composed of primer F1 and a different reverse primer R1 downstream of the CLCC1 start codon, is expected to produce a ~1000 bp PCR product if any allele maintains the authentic 5′ region of the gene. This is observed for the HeLa KO cell line, indicating an incomplete knockout of likely one CLCC1 allele.

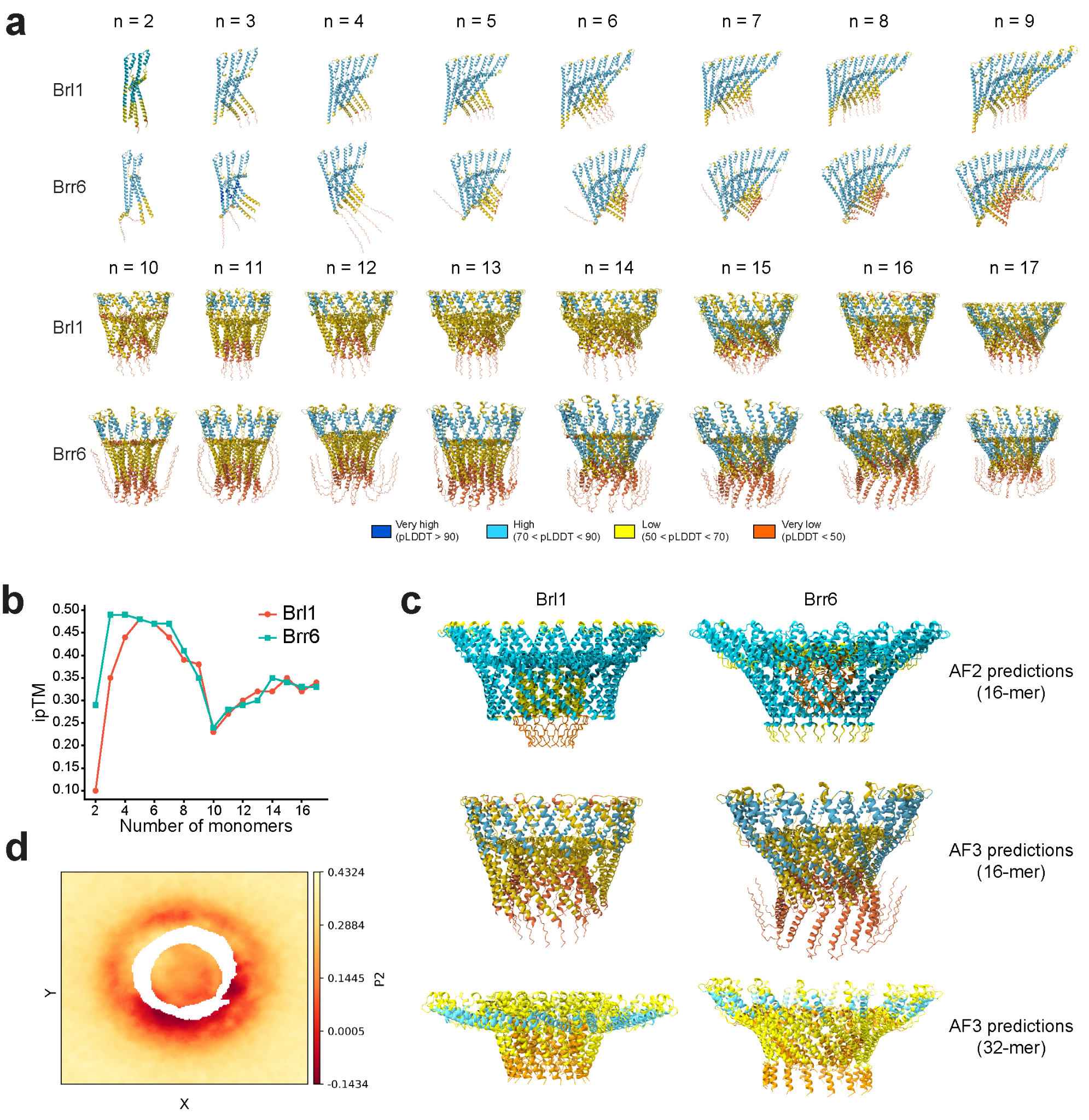

**Supplemental Figure S5: AlphaFold-predicted homo-oligomers of Brl1 and Brr6. a)** AlphaFold3 predictions of Brl1 and Brr6 complexes for 2-17 subunits (n) colored by the residue-specific confidence score (pLDDT). **b)** Interchain predicted template modeling (iPTM) of the predicted complexes in (a). **c)** Prediction of Brl1 and Brr6 16-mer from AlphaFold2 (top), AlphaFold3 (center) and Brl1/Brr6 16-mer from the 32-mer prediction of Brl1:Brr6 (16:16) with AlphaFold3 (bottom) colored by the residue-specific confidence score (pLDDT). **d)** Rank 2 lipid order parameter (P2) of lipid tails from coarse-grained simulations of the Brl1 16-mer in a lipid bilayer (Fig 3d).

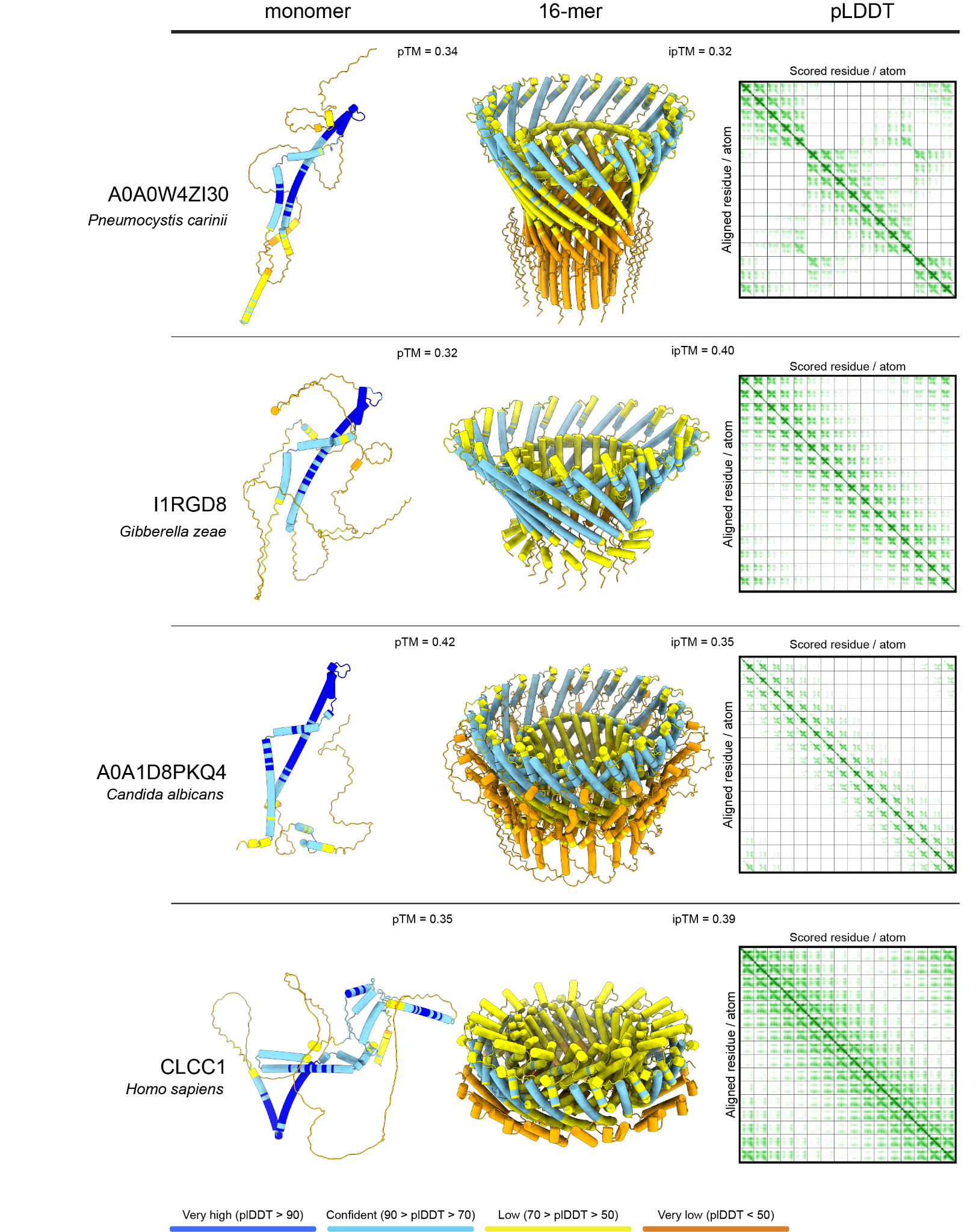

**Supplemental Figure S6: AlphaFold3 predicts homo-multimeric complexes for different Brl1/Brr6 homologues and CLCC1.** Monomers and 16-mers of the indicated *BRL1/BRR6* homologues and CLCC1 colored by residue specific confidence score (pLDDT) and their respective pLDDT plot. The unstructured C-terminus of CLCC1 was truncated in this prediction, and only residues 1–365 are included.

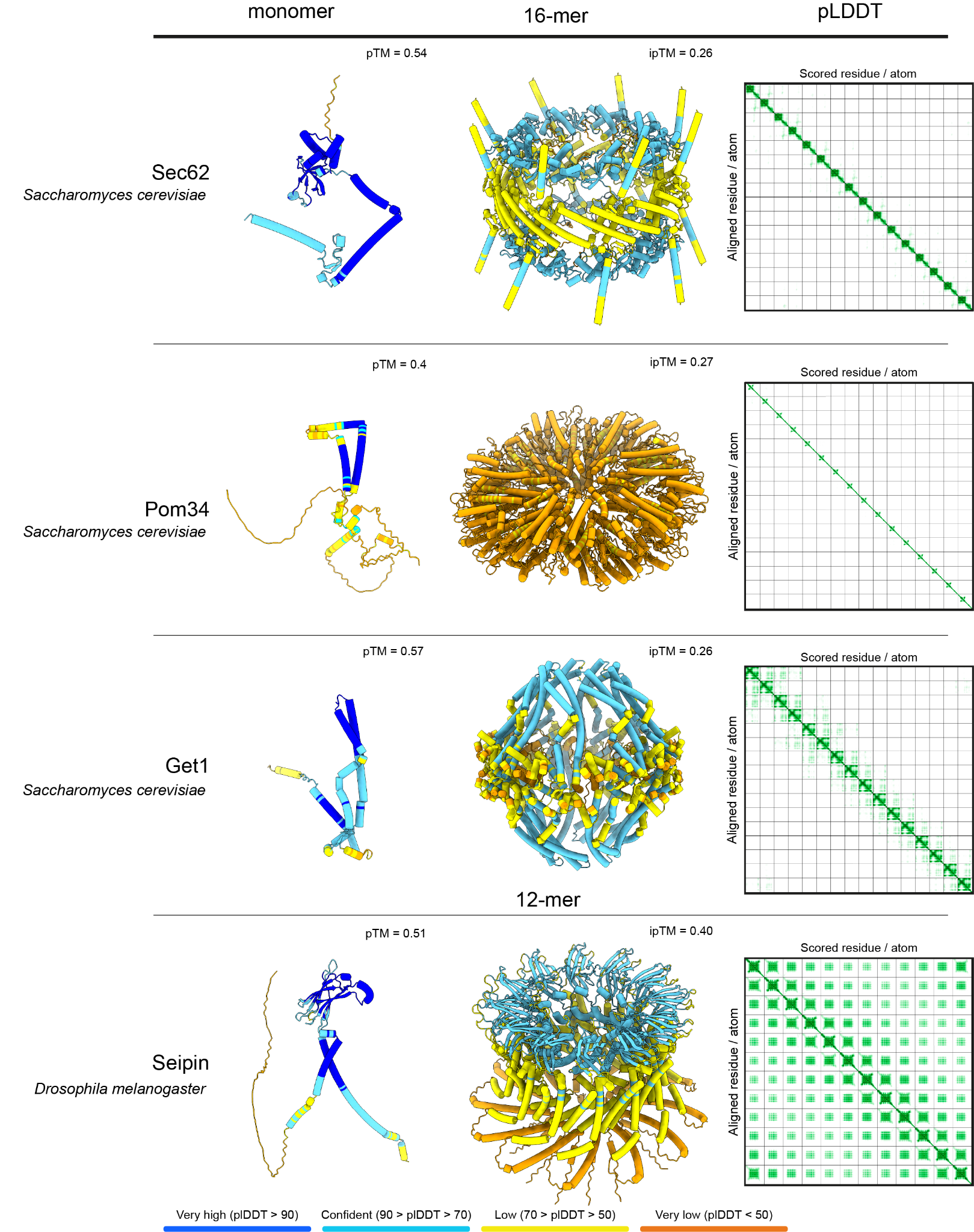

**Supplemental Figure S7: AlphaFold3 predictions for randomly chosen two- or three-pass α-helical transmembrane proteins.** Monomers and 16-mers of randomly chosen *S. cerevisiae* transmembrane proteins colored by residue-specific confidence score (pLDDT) and their respective pLDDT plot. *Drosophila melanogaster* Seipin is known to form a 12-mer^92^.

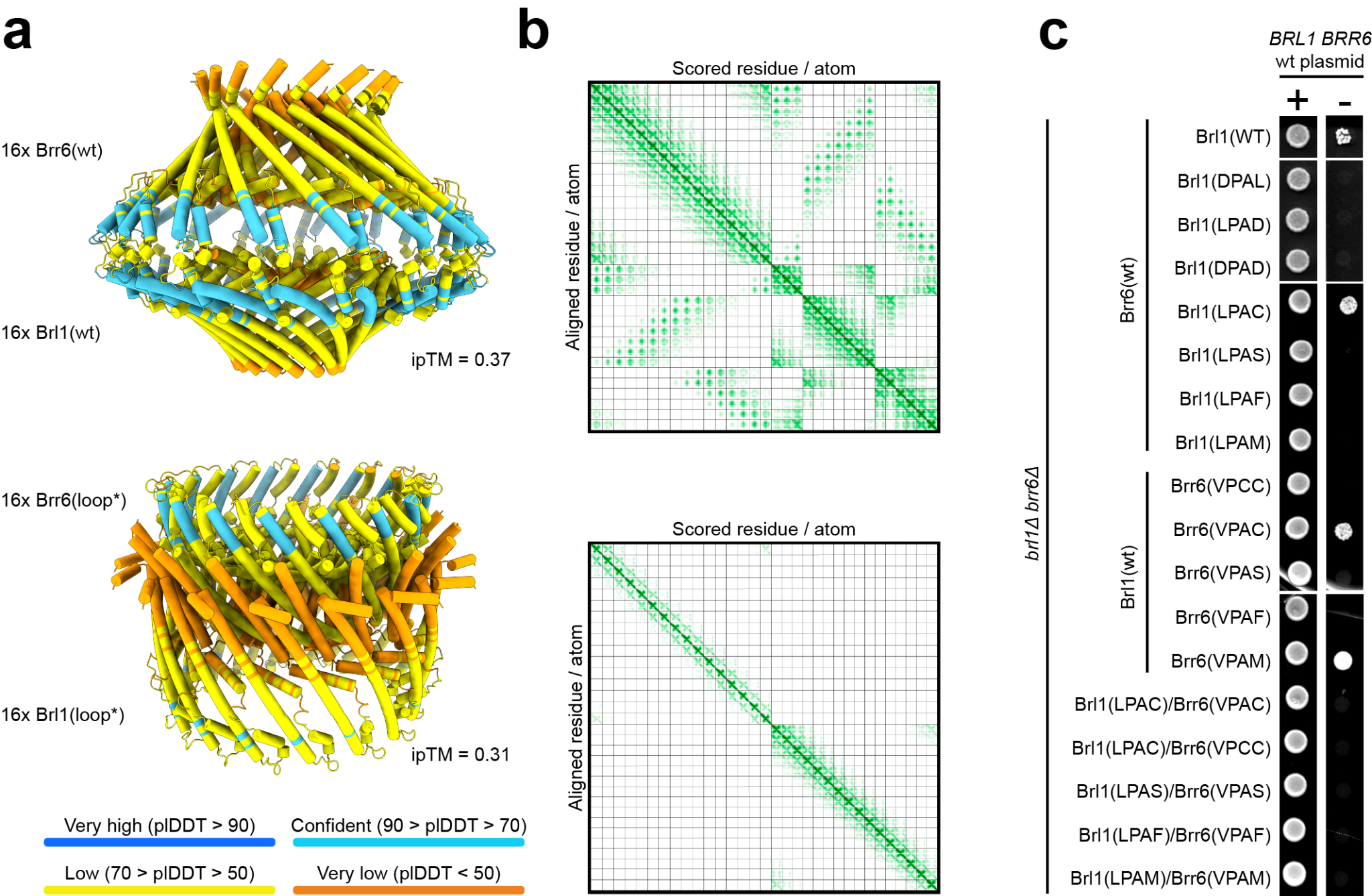

**Supplemental Figure S8: Disruption of the hydrophobic loop motif renders Brl1 and Brr6 non-functional.** **a**) AlphaFold3 prediction for 16 copies of Brl1 and 16 copies of Brr6 with an unperturbed interaction motif (top) and mutated hydrophobic loops (bottom) colored by the pLDDT. **b**) Predicted aligned error plots for the models in (a). **c**) Viability of cells expressing Brl1 and Brr6 hydrophobic loop variants from the endogenous promoter using the plasmid shuffle assay in the presence or absence of a *BRL1*/*BRR6* wt plasmid.

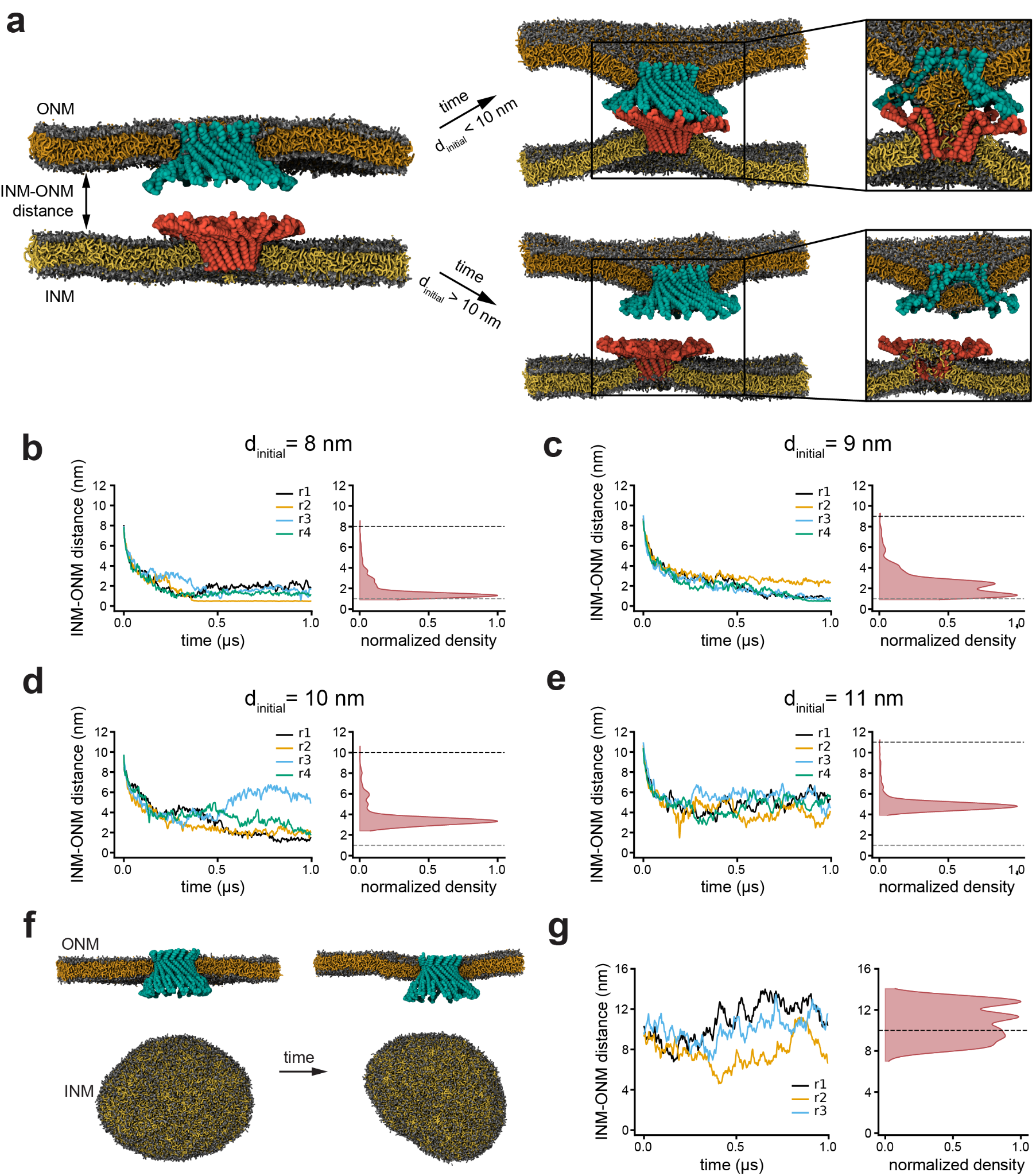

**Supplemental Figure S9: An initial distance d_initial_ of < 10 nm is required for Brl1-Brr6 interaction.** **a)** Coarse-grained MD simulation setup featuring two opposing lipid bilayers at an initial minimum distance (d_initial_) with Brl1 and Brr6 16-mers inserted into the INM and ONM, respectively (left). The right panel shows representative final simulation snapshots illustrating two distinct outcomes: lipid mixing between membranes for d_initial_ < 10 nm and no lipid mixing for d_initial_ > 10 nm. **b-e)** Minimum intermembrane (INM-ONM) distance during the simulation for systems at multiple initial minimum distances (d_initial_ = 8, 9, 10, 11 nm). **f)** Coarse-grained MD simulation of a planar lipid bilayer with a Brr6 16-mer inserted, opposed to a vesicle with no protein. **g)** Minimum intermembrane (INM-ONM) distance during the simulation in f). INM – inner nuclear membrane, ONM – outer nuclear membrane.
